# Can’t carve air, can’t weave names: Plant species used in Colombian artisan crafts are not equally assessed for vulnerability at the international and national level

**DOI:** 10.1101/2024.01.24.573691

**Authors:** Katherine Victoria Hernandez, Federico Andrade-Rivas, Felipe Zapata, Natasha Batista, Anaid Cárdenas-Navarrete, Armando Dávila Arenas, Guido A. Herrera-R, Kelley E. Langhans, Dallas Levey, Andrew Neill, Oliver Nguyen, Natalia Ocampo-Peñuela, Sergio Sánchez Lopez, Alejandra Echeverri

**Affiliations:** Institute of the Environment and Sustainability, University of California Los Angeles, Los Angeles, CA, USA; School of Public Health and Social Policy, University of Victoria, Victoria, BC, Canada; Instituto de Salud y Ambiente, Universidad El Bosque, Bogotá, Colombia; Department of Ecology and Evolutionary Biology and Center for Tropical Research, Institute of the Environment and Sustainability, University of California Los Angeles, Los Angeles, CA, US; The Natural Capital Project, Stanford University, Stanford, CA, USA; Department of Integrative Biology, University of California Berkeley, Berkeley, CA, USA; Independent Musician, Bogotá, Colombia; Department of Ecology and Evolutionary Biology, The University of Tennessee, Knoxville, TN, USA; Department of Fish and Wildlife Conservation, Virginia Tech, Blacksburg, VA, USA; Department of Ecology and Evolutionary Biology, Stanford University, Stanford, CA, USA; Department of Botany, Trinity College, Dublin, Ireland; Department of Environmental Studies, University of California Santa Cruz, Santa Cruz, CA, USA; Doerr Sustainability School, Stanford University, Stanford, CA, USA; Department of Environmental Sciences, Policy, and Management, University of California Berkeley, Berkeley, CA, USA

**Keywords:** Biocultural diversity, Biodiversity conservation, Ethnobotany, Indigenous Peoples and Local Communities, Plants, Colombia

## Abstract

Humanity has maintained cultural connections with our environments for time immemorial. Plants and artisan crafts are a prime example, as craft purpose, skill, design, and species used can vary greatly between communities and the loss of a critical plant species can result in a loss of access to cultural craft practices. To mitigate global biodiversity loss, conservationists are faced with the challenge of assessing species vulnerability to extinction and prioritizing species for conservation funding using information instruments, like the IUCN red list. This process does not necessarily consider a species’ cultural importance. In this paper we sought to address this gap for plant species used in artisan crafts in Colombia. We aim to answer the following: (1) how represented are endemic species in artisan crafts; (2) how threatened are artisan craft species according to (a) international and (b) national vulnerability status? We used the number of species-associated common names as a proxy for cultural awareness. We found that continentally regional species were far more represented in Colombian artisan crafts than national endemics. We also found a strong positive relationship between number of common names and national vulnerability assessment status, but no statistically significant relationship for international vulnerability status. Based on our results, well-known plants used in Colombian artisan crafts are more likely to be assessed nationally than internationally. While the IUCN is thorough in their recommendations, more can be done to prioritize the inclusion of conservation assessments for species based on their contributions to cultural diversity.

**Positionality statement:** We are 14 scientists and practitioners who are deeply committed to the conservation of nature and culture in a changing world. We are trained in diverse fields including ecology, evolutionary biology, botany, music, anthropology, law, and public health. We all have postgraduate academic education (Masters or PhDs underway) but most of us are early career scholars. Six of us grew up in Colombia and we represent many places including Mexico, the United States, Ireland, Chile, Brazil, Germany, and Viet Nam. None of us self-identify as Indigenous.

## 1. Introduction

Amidst an unprecedented biodiversity crisis lies an intertwined loss of cultural identity and diversity (Pretty et al., 2009). Not all peoples are equally affected. Ceremonies, customs, and traditions of Indigenous Peoples and Local Communities (IPLC) around the world depend on the reciprocal relationships between humans and other species, including some on the brink of extinction (Hill et al., 2020; Nabhan, 2000; Turner et al., 2000). In 1996, the first ‘Endangered Languages, Endangered Knowledge, Endangered Environments’ working group gathered in Berkeley, California (US) and discussed the shared threats to and vulnerability of global linguistic, cultural, and biological diversity (Maffi & Woodley, 2012, p.xix-xxi). The intersection of these three diversities, otherwise known as biocultural diversity, has grown as a conservation topic of interest in the years since (Maffi & Woodley, 2012, p. xix-xxi).

Limited funds and time necessitate prioritizing species for conservation (Blackman et al., 2014; Arponen, 2012). The International Union for Conservation of Nature’s (IUCN) Red List of Threatened Species is a comprehensive database of species so far assessed for global extinction risk, a critical criterion considered in species conservation prioritization (IUCN, 2023; Arponen 2012). Conservation planning, however, tends to occur at local, national, and regional scales, where applying global vulnerability assessment results is not always practical. Numerous national red lists have been published, many of which have used IUCN Criteria systems for their assessments (IUCN, 2023).

In addition to extinction risk, it is common practice to prioritize species by ecological value and potential impact of loss (Cadotte, Carscadden, & Mirotchnick, 2011; Chapin Iii et al., 2000; Rodriquez et al., 2015; Witting, Tomiuk, & Loeschcke, 2000). Ecological keystone species (EKS) are species with such disproportionately large impacts on their community or ecosystem, relative to their abundance, that their absence may lead to a major loss of ecological function and overall health (Power et al. 1996). Species that fit the EKS model are thus often proposed as priority species. Similarly, the term “cultural keystone species (CKS)” was originally coined to describe “culturally salient species that shape in a major way the cultural identity of a people, as reflected in the fundamental roles these species have in diet, materials, medicine, and/or spiritual practices” (Garibaldi & Turner, 2004). For example, *kalo* (taro, *Colocasia esculenta)* is a primary staple in Hawaiian Indigenous cuisine (Winter, Lincoln, & Berkes, 2018), Black ash (*Fraxinus nigra*) has medicinal and ceremonial importance for multiple Tribal Nations in North America (Siegert et al., 2023), and guassa grass (*Festuca macrophylla*) has many uses for communities living in the northern highlands of Ethiopia, including food, clothes, and building material (Chengere et al., 2022). CKS has a similarly narrow definition to EKS, as it looks specifically at species whose loss would lead to major cultural changes and whose initial absence would likely have resulted in major differences in the resulting culture(s), like the functioning of economy, food security, and utility (Garibaldi & Turner, 2004). *Culturally important species* (CIS) hold less distinct roles than CKS do, but are widely recognized in a culture and widely used (Reyes-García et al., 2023). Previous work has elucidated CIS vulnerability by developing a biocultural vulnerability index (Appendix Fig. S.1.), which uses IUCN red-list status to determine species’ biological vulnerability and uses language vulnerability as a proxy for cultural vulnerability (Reyes-García et al., 2023).

Why some species are culturally important over others is highly variable across cultures, space and time (Platten & Henfrey, 2009; Bruyere, Trimarco, & Lemungesi, 2016; Bond et al., 2019). Hand-made products made over generations—artisan crafts, as we refer to them in this paper—have been developed in the context of materials available, much of which are plant species (Balick & Cox, 1996, pages 1-8). Basketry across the American continents is a prime example of this. Basket traditions vary across peoples, each with their own craft practices for weaving, using different plant species as raw material, making baskets at different times and by different individuals, as well as applying various patterns and decorations. In this way, basketry and other forms of artisan craftsmanship are unique to each cultural group and are the physical representations of unique human-plant relationships (Salmón, 2000).

Plants are less prioritized for conservation funding compared to animal species (Balding & Williams, 2016; Goettsch et al., 2015). The most visible plant species are typically those used for food, fiber, and ornamental use, and those with long existing cultural significance, often tied to identity, empathy, and other deeply held interactions (Balding & Williams, 2016). The more culturally visible a species is, the better chance it has to be funded for conservation and recovery (Balding & Williams, 2016). This could suggest that bioculturally important plant species are likely already being targeted for conservation. However, there is a strong North American and European bias in ecological research, both for where studies are done and by whom (Martin, Blossey, & Ellis, 2012; Di Marco et al., 2017). Geographic biases are reflected in global vulnerability assessments, like the IUCN Red List, and can skew where conservation funding is directed (de los Rios, Watson, & Butt, 2018). National species assessments may be better at including CIS within national borders, but even so, local communities may not have the power, resources, or representation to ensure protection for CIS.

We approach this issue with the following questions: (1) how represented are endemic plant species in Colombian artisan crafts; (2) how threatened are endemic artisan plant species used in Colombian artisan crafts according to (a) international vulnerability status, and (b) national vulnerability status; and (3) what relationship, if any, exists between artisan plant species’ (a) international and (b) national vulnerability status and how visible the species is, with number of common names as a language proxy for the present cultural awareness for a species?

## 2. Materials and Methods

### 2.1. Case study selection

Colombia is one of the world’s four most bioculturally diverse countries, established by a countries’ number of languages, religions, and ethnic groups present per country, as well as number of bird, mammal, and plant species (Loh & Harmon, 2005; Andrade, 2011; Arbeláez-Cortés, 2013). Colombia is ranked first in bird species richness, second in amphibians, freshwater fish and butterflies, third in reptiles, fourth in plant species, and fifth in mammals (Andrade, 2011; Rodríguez-Zapata & Ruiz-Agudelo, 2021). Over 52 million people live in Colombia, across 32 governmental departments (DANE, 2018). Colombia is home to >170 ethnic groups (including Indigenous, Black, Afros, Raizales, Palenqueros, Gitanos, and Rrom) who steward and have legal right over ∼24% of the country’s territory (DANE, 2018). Colombia’s social, political, and ecological history have led to a nation with an extremely high level of biodiversity and a complex web of relationships between people and the environment, all of which are threatened by anthropogenic activities (e.g. mining, forest clearing, pollution, climate change) (Ocampo-Peñuela et al., 2022; Blackmen et al., 2014).

While Colombia is ranked fourth in plant species richness, it is second in plant biodiversity (Diazgranados et al.; 2022b). Through the efforts of the Useful Plants and Fungi of Colombia (UPFC) project, over 36,000 Colombian plant and fungi species have been identified and catalogued, encompassing 5,346 genera and 843 families (Diazgranados et al., 2022b). At least 7,790 of Colombian’s plant species are useful to humans (Diazgranados et al., 2022b; Bernal & Sánchez, 2022; Gori et al., 2022). 2363 plants identified as material-providing species (Diazgranados, 2022). Like edible and medicinal plants, material species hold cultural importance, but fewer studies have focused on material plant species as a threatened group of cultural importance.

We focused on plant species for their high representation in artisan crafts, like basketry, and the recognized importance of plant species in other South American nations (Bernal et al., 2013; Santamaría, 2017; Vargas & van Andel, 2005; Cordoba Arroyo, 2018; Reyes-García, 2006; Balick & Cox, 1996; García et al., 2013).

### 2.2. Data collection

We searched four data sources to identify species: Artesanías de Colombia; Universidad Nacional de Colombia – Catálogo: Nombres Comunes; Royal Botanical Gardens KEW - Plants of the World Online (POWO); and the Catalogue of Useful Plants of Colombia (*Catálogo de Plantas Útiles de Colombia*) (Table S.1.).

We performed an iterative search process encompassing five rounds of data collection to build a database of plant species.

The first two iterations were done by reviewing the *Artesanías de Colombia* website (Artesanias de Colombia, 2023). *Artesanías de Colombia* is a catalog of crafts from 1000 businesses across Colombia, sold between 1990-2020, indicating their origin and primary plant species used. We collected the common and scientific names of primary plant species when found, and the type of crafts they were used for. Given that not all plant species had associated documents with the information we needed (i.e. full species name, all common names, departments used or cultivated in, plant family, vulnerability status), we did a third iteration by searching the *Universidad Nacional de Colombia, Nombres Comunes* database. The Nombres Comunes database contains more than 18,000 Spanish language common names used to designate plants in Colombia, and some names in Creole English as used in the Caribbean islands of San Andrés and Old Providence (Bernal et al., 2017b). Common names from Indigenous languages were likely underrepresented in both the Artesanías de Colombia archive and Nombres Comunes database, and so underrepresented in our data (See “Discussion 4d.: Limitations”).

On the fourth iteration we reviewed the *Catálogo de Plantas y Líquenes de Colombia* (Bernal, Gradstein, & Celis, 2019). This is an inventory of the approximately 28,000 plant species identified in Colombia. Each plant species profile contains available information on plant size, ecological and political regions where species is or has been present, current conservation status, and global distribution, (López & Prada, 2014). Fifth, we searched *the Catalogue of Useful Plants of Colombia* and POWO for species taxonomy and distribution (Negrão et al., 2022). Last, we reviewed the IUCN Red List of Threatened Species and added information on the vulnerability status for each species (IUCN, 2022). We listed plants native only to Colombia as “endemic.” Species native in Colombia and at least one other neighboring country were listed as “regional.” All other species were labeled as “introduced.” Three sources were consulted to evaluate endemic and vulnerability status: Plants of the World database for endemic status, the Catálogo de Plantas y Líquenes de Colombia for nationally documented vulnerability status, and the IUCN Red List for international vulnerability status and categories (Table S.1.).

We built a database and collected information on the taxonomy of species by recording species names, scientific names, taxonomic family, local name(s), other names, taxon, IUCN status, departamento(s) (i.e., second tier of administrative governance, equivalent to states or provinces) of Colombia where the species is used for artisan crafts, and how they are used (e.g., woodworking, basketry, natural dyes).

### 1.3 Taxonomic Validation

We corrected for synonymous names (common and scientific) and unverifiable references. Species names were found through Artesanías de Colombia documents through common names, photographs, geographic identifiers, and other additional information. When a species name was not directly listed in archival documents, or cases of reasonable doubt (i.e. misspelling, multiple scientific names attributed to single plant) species names were cross-checked with the Catalogue of Useful Plants of Colombia, and Plants of the World Online (POWO) web repository to determine likely plant species identity (See Table S1). Plant species that were listed in archives but not in the Catalogue of Useful Plants of Colombia were heavily cross-researched before deciding to include or omit from the final database. We only included plants that could be tracked down to the species level.

### 1.4. Common Names as a Proxy for Cultural Awareness

Over 65 Indigenous languages are spoken in Colombia, in addition to two creole languages, Romani, Spanish, and more (*Plan Decenal de Lenguas Nativas de Colombia 2022-2032,* 2022). The social-political history of Colombia is such that traditional craft artisans may be learning and sharing their craft in multiple languages, and in languages not connected to the craft’s cultural origins. Multiple cultural groups may share a language while not sharing the same craft practices.

We use local common names as proxy for the cultural awareness of a plant species in Colombia beyond the interests of the international scientific community, where every recognized species is assigned a Latin name by rule. We assume that if a plant species has many common names, it is the result of many people, over time, talking about said plant to each other, to people of other cultural groups, in different languages and in different places. A plant species with multiple common names and name spellings may be well-known by many, while plants with few common names may only be known by a few. Different names may also be assigned to different useable parts of a plant (i.e. corn cob, husk, stalk). Some plants may have the same common name shared across many cultural groups, but over time gain discrepancies in spelling; thus, we count different spellings of very similar names as separate names.

### 1.5. Statistical Analysis

We used R version 2023.03.1+446 as our programming language to perform statistical analyses (R Development Software team, 2023). We analyzed five variables: species identity, common names, endemic status, national vulnerability status, and international vulnerability status. We created two frequency tables: (1) the number of species with each endemic status, and (2) the number of local common names for each species. Boxplots and QQ-plots were created to check for outliers and normal distribution in the data. We used a p-value of 0.05 for all tests.

#### 1.5a. Inquiry 1: Endemic status of artisan plant species

For our first inquiry we asked how many local common names were associated with national endemic artisan plant species in comparison to non-endemics? We hypothesized that at least one of the endemic status categories would have a different median for the number of local names associated with each species. Our null hypothesis was that the median number of local names across the categories of endemic status would be the same. To test this hypothesis, we performed Kruskal-Wallis tests, as we were testing one categorical independent variable (endemic status) versus one numeric continuous (number of local common names) response variable that is not normally distributed.

#### 1.5b. Inquiry 2: Vulnerability of endemic artisan plant species

For our second inquiry we asked are endemic artisan plants more likely to be endangered according to (a) international, and (b) national vulnerability statuses? We hypothesized (a) there is an association between IUCN vulnerability status and endemic status, and (b) there is an association between Colombian vulnerability status and endemic status. Our null hypotheses were (a) no association between IUCN vulnerability and endemic status, and (b) no association between national vulnerability status and endemic status. To test these hypotheses, we performed Chi-squared tests (when observations are >5) and Fisher’s exact tests (when observations <5 or 0).

#### 1.5c. Inquiry 3: Vulnerability of artisan plant species across endemic status

For our last inquiry we asked what relationship, if any, exists between an artisan plant species’ international and national vulnerability status and the cultural awareness for the species, according to (a) the IUCN Red List, an international vulnerability assessment, and (b) the Universidad Nacional de Colombia, Catálogo de Plantas y Líquenes de Colombia, where the reported species vulnerability statuses were assessed nationally. We hypothesized (a) at least one of the IUCN categories has a different median of the number of local names used to call each plant; and (b) at least one of the national vulnerability categories has a different median of the number of local names used to call each plant. Our null hypotheses were that the median number of local names across the categories of (a) IUCN status, and (b) national vulnerability status, were the same. To test this hypothesis we performed Kruskal-Wallis tests, with Dunn’s post-hoc tests for significant results.

## 3. Results

We found 103 references to plants used for making crafts in the form of common and scientific names, prior to checking names for redundancy and taxonomic validation. We found 76 distinct plant species: two endemics (*Astrocaryum malybo* and *Saurauia ursina*), 15 introduced, and 59 regionals (Appendix, Table S2). Results showed a median 10.0 and a standard deviation of 11.6 local common names per species across all categories. Common names were not normally distributed across endemic status (Appendix – Plot S.1.). Three outliers were identified: *Crescentia cujete* (45 common names), *Lagenaria siceraria* (50 common names), and *Zea mays* (52 names); also known as the calabash tree, the bottle gourd, and corn respectively. The calabash tree is regional, while the bottle gourd and corn are introduced species and internationally distributed. When removed, summary statistics showed that outliers skewed data slightly (Median = 9.0, SD = 9.006952).

### 3a. Number of Local Common Names for Endemics

There were no significant differences across endemic status in terms of the number of local common names, with the outliers (Kruskal-Wallis chi-squared: H(2) = 2.0159, p = 0.365) or without (Kruskal-Wallis chi-squared: H(2) = 5.4366, p = 0.06599) (Table 2) (Figure 2). We cannot reject the null hypothesis in both cases, and so our results show that nationally endemic artisan plants do not have more local common names than regional or introduced artisan plant species. We performed the same analysis after the endemic category was removed, which had a small sample size (n=2), and found the same conclusions (Kruskal-Wallis chi-squared: H(1) = 1.6616, p-value = 0.1974).

**Figure 1.**
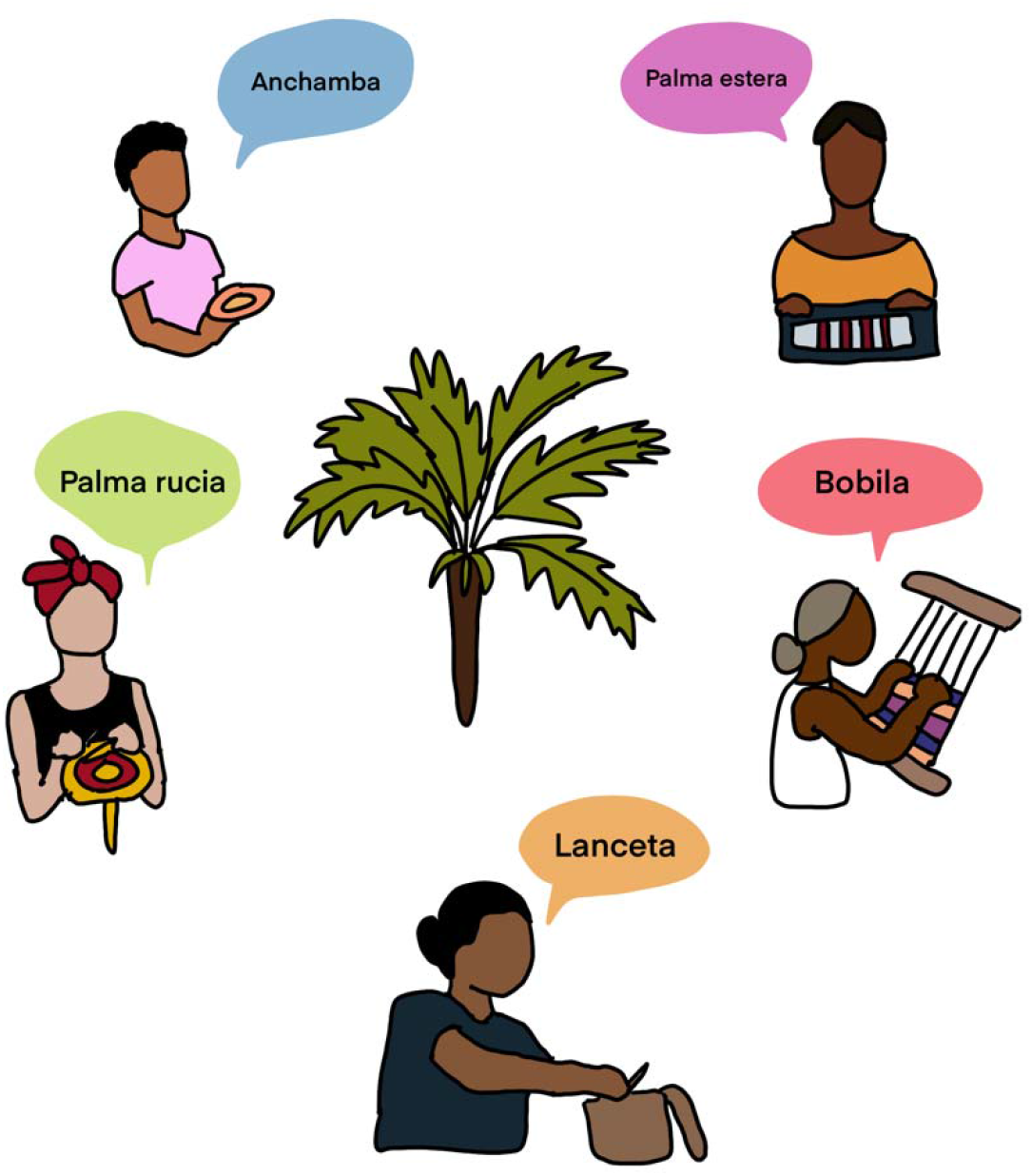
[Color preferred for print]. Conceptual diagram showing how a plant species (*Astrocaryum malybo*) can be used and depended on for a number of different craft practices and be called by different names across uses and communities.

**Figure 2.**
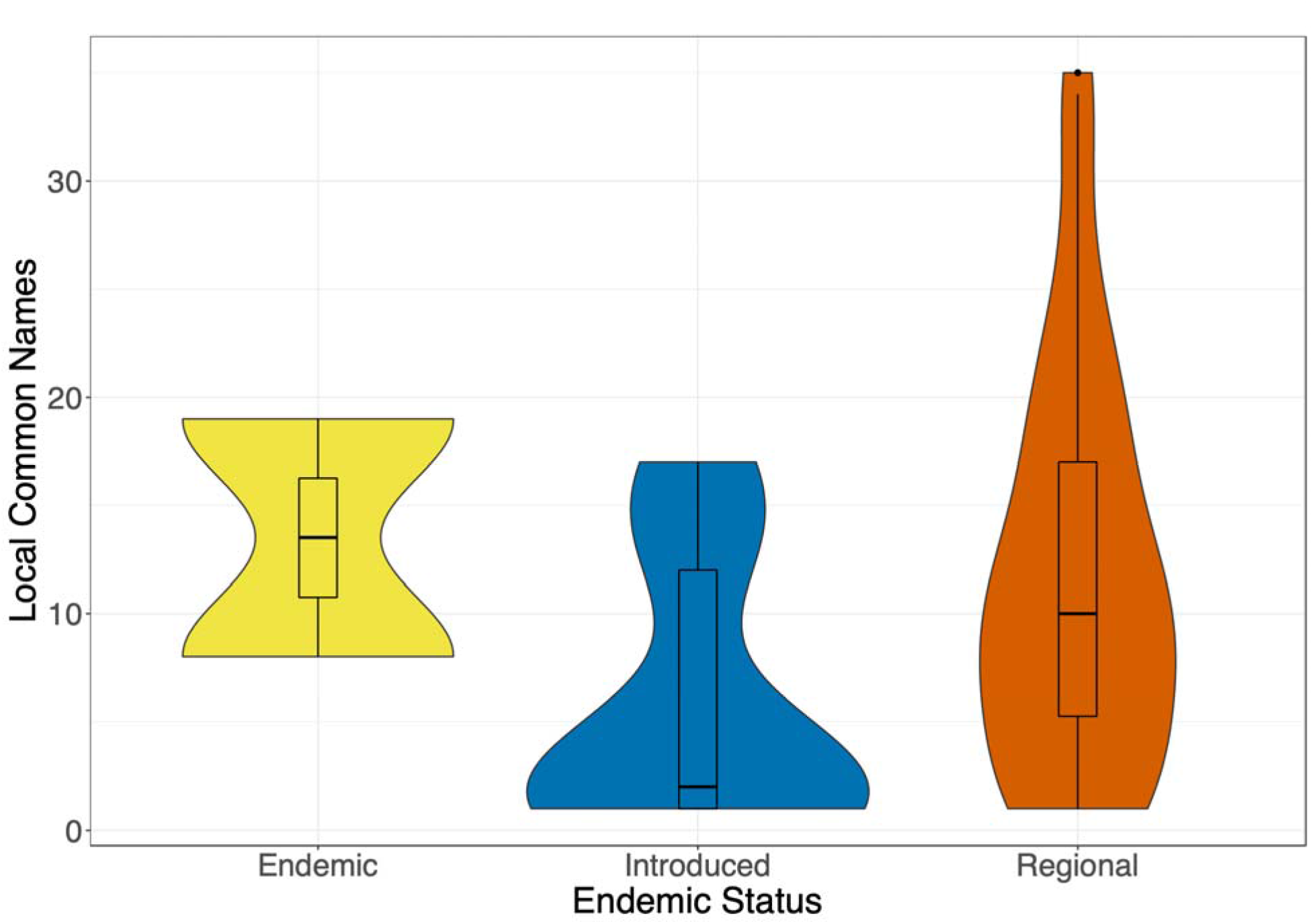
[Color preferred for print]. Number of local common names per species by endemic, introduced, and regional status groups. Outliers (3) have been removed.

### 3b. Vulnerability of Endemics

Of the nine potential IUCN categories possible for our analysis (“Unknown”, “Data Deficient”, “Least Concern”, “Near Threatened”, “Vulnerable”, “Endangered”, “Critically Endangered”, “Extinct in the Wild”, “Extinct”) only five were represented in our data set (“Unknown”, “Data Deficient”, “Least Concern”, “Vulnerable”, and “Endangered”) and the “Data Deficient” and “Endangered” categories each had only one data point (Bland et al., 2017).

A contingency table of endemic status versus IUCN status (Table 1) was zero inflated, so we performed a Fisher’s Exact Test for Count Data with simulated p-value (based on 2000 replicates)(Table S.2.). Results were insignificant (p-value = 0.6847), thus we cannot reject the null hypothesis (Table S.2.). The endemic species category contained multiple zeros and was removed before running the test again. Results remained insignificant after removal (p-value = 0.8586). We have no evidence to support that endemic artisan plants are more endangered, or an association between international vulnerability status and endemicity.

**Table 1.** Contingency table of endemic status vs. IUCN international vulnerability status and ColPlant A national vulnerability status.

| IUCN Status | Endemic status |  |  |
| --- | --- | --- | --- |
|  | Endemic | Introduced | Regional |
| UN | 2 | 8 | 24 |
| DD | 0 | 0 | 1 |
| NT | 0 | 0 | 0 |
| LC | 0 | 7 | 30 |
| VU | 0 | 0 | 3 |
| EN | 0 | 0 | 1 |
| ColPlant A status | Endemic | Introduced | Regional |
| EN | 1 | 0 | 2 |
| LC | 0 | 3 | 15 |
| NE | 1 | 6 | 33 |
| NT | 0 | 0 | 4 |
| UN | 0 | 4 | 2 |
| VU | 0 | 0 | 2 |

Results were similar at the national level. The contingency table for endemic status and national vulnerability status (Table 1) was also zero inflated, and another Fisher’s Exact Test for Count Data was run with simulated p-value (based on 2000 replicates). There was no significant relationship before excluding endemic species (p-value = 0.08646) or after (p-value = 0.1419). We have no evidence to support the hypothesis that endemic plants are more endangered, or an association between national vulnerability status and endemicity.

A qualitative inspection of the violin plots comparing IUCN and national vulnerability statuses and number of local names (Fig 2) show notably large ranges for common names associated with Unknown (UN) and Least Concern (LC) species in IUCN evaluations, and for common names associated with Not Evaluated (NE) and LC species in national vulnerability assessments. The national vulnerability status and number of local names violin plot illustrate the positive relationship between the two variables. The median number of local names for species with no profile (UN) (Med = 1.5) or not evaluated (NE) (Med. = 7.00) were below the median of all local names (Med. = 9.00). The highest represented vulnerability statuses, Vulnerable (VU) and Endangered (EN), both had medians well above the median for all local names (Med. = 17.50; Med. = 18.00). Near Threatened (NT) species had a median (Med= 8.5) close to the median of NE species and the overall median. The median number of local common names for LC species (Med. = 14.50) were comparable to VU and EN. Not Evaluated (NE) and Least Concern (LC) plant species had the largest ranges (35 and 29, respectively) in contrast to the average of all others (11.25).

### 3c. Vulnerability and Cultural Awareness

There were no significant differences in the number of common names across IUCN status with outliers (Kruskal-Wallis chi-squared: H(4) = 3.8118, p-value = 0.4321) or without (Kruskal-Wallis chi-squared: H(4) = 4.068, p-value = 0.3969). Our results show that artisan plants classified as endangered by the IUCN do not have more local names than other categories under the same assessment criteria. We performed the same analyses by excluding the DD and EN categories, which had a small sample size (n=1 each), and found the same conclusions (Kruskal-Wallis chi-squared: H(2) = 1.4874, p-value = 0.4754) (Table 2).

**Table 2.** Kruskal-Wallis results.

| Independent variable | Response variable | Data | Kruskal-Wallis Chi-squared | Degrees of freedom | P value |
| --- | --- | --- | --- | --- | --- |
| Endemic status | Count of local names | All data | 2.02 | 2 | 0.365 |
|  |  | Without outliers | 5.43 | 2 | 0.06 |
|  |  | Excluding endemic category | 1.6616 | 1 | 0.1974 |
| IUCN status | Count of local names | All data | 3.8118 | 4 | 0.4321 |
|  |  | Without outliers | 4.0868 | 4 | 0.3969 |
|  |  | Excluding DD and EN categories (each n=1) | 1.4874 | 2 | 0.4754 |
| National vulnerability status | Count of local names | All data | 17.848 | 5 | 0.00314 |
|  |  | Without outliers | 16.548 | 5 | 0.00544 |

However, there were significant differences between national vulnerability status and number of common names, with outliers (Kruskal-Wallis chi-squared: H(5) = 17.848, p = 0.003143) and without outliers (Kruskal-Wallis chi-squared: H(5) = 16.548, p = 0.005441) (Table 2) (Figure 3). There is evidence to support that at least one of the medians is different from the others, meaning at least one of the categories of national vulnerability status has more or less local names than expected by chance. A Dunn’s post-hoc test was performed to see which category’s median differed from all others (Table 3). Results show that LC-NE are different, and that LC-UN are different, all others the same. This shows that plants assessed as Least Concern, nationally, have more local names than Near Endangered or Unknown plants.

**Figure 3.**
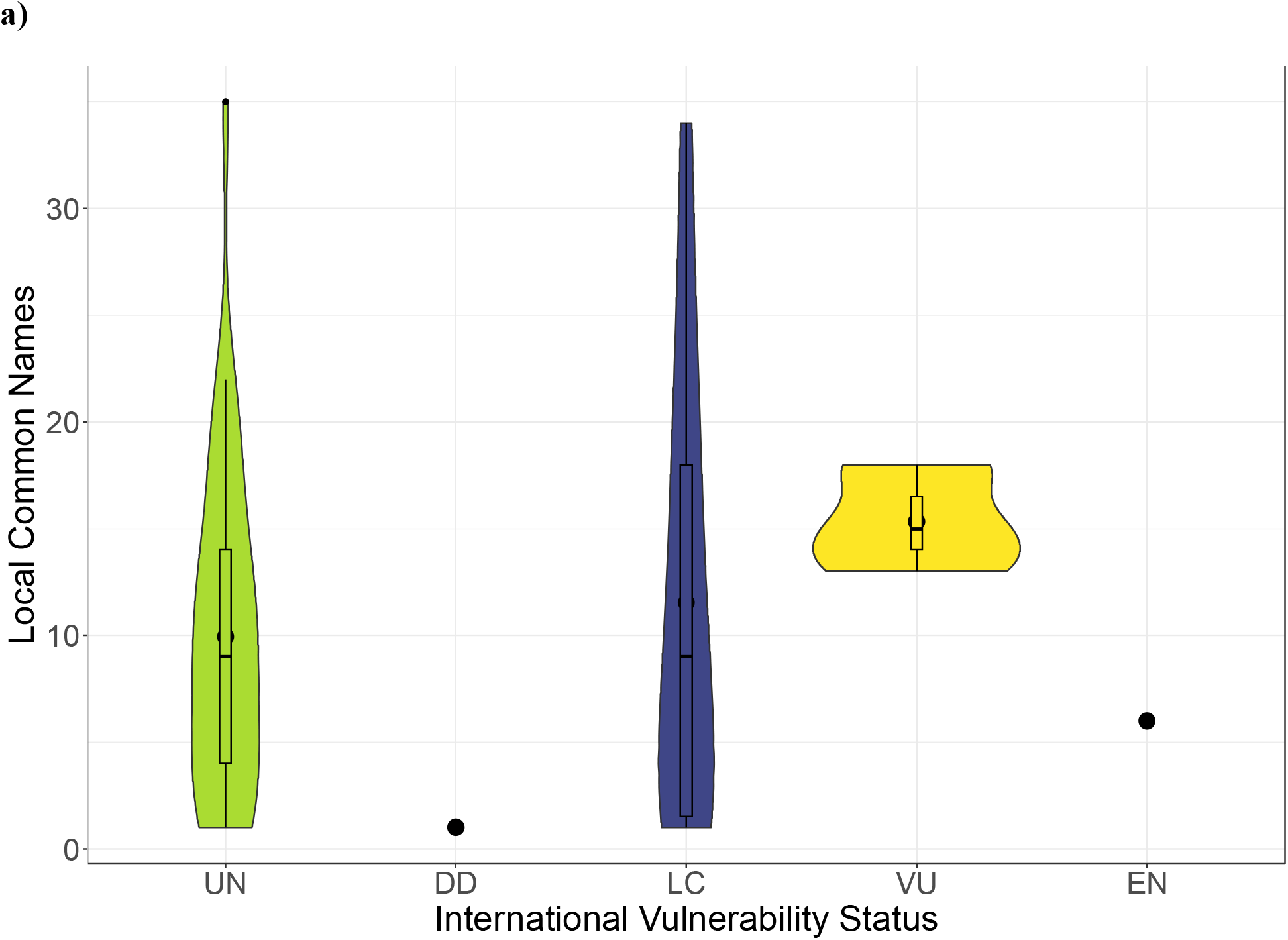
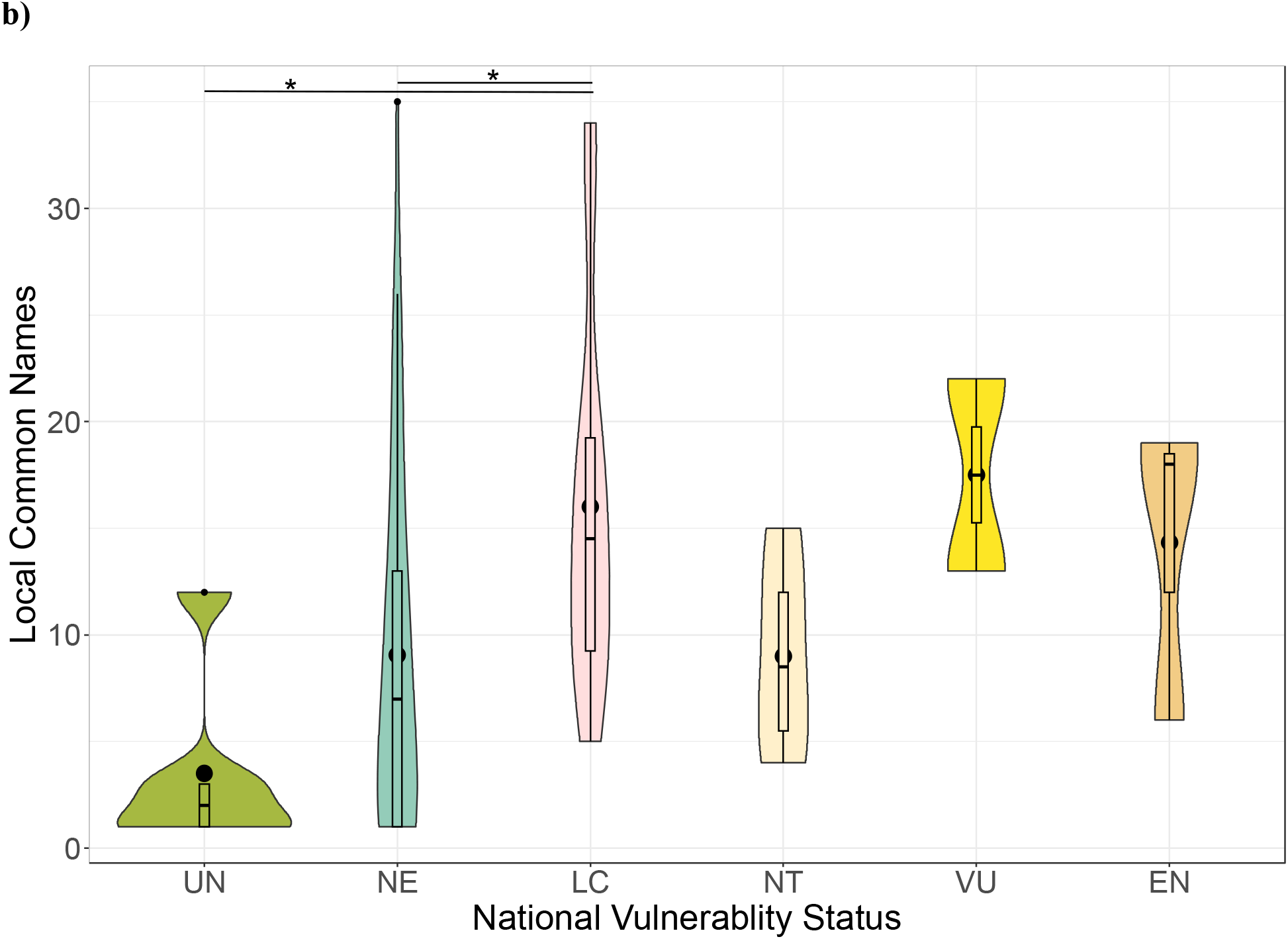
[Color preferred for print]. Number of local common names by **a)** international vulnerability status (statuses possible: Unknown (UN), Data Deficient (DD), Least Concern (LC), Near Threatened (NT), Vulnerable (VU), Endangered (EN), Critically Endangered (CE), Extinct in the Wild (EW), Extinct (EX)) as assessed by the International Union for Conservation of Nature (IUCN), and **b)** national vulnerability (statuses possible: Unknown (UN), Not Evaluated (NE), Least Concern (LC), Near Threatened (NT), Vulnerable (VU), Endangered (EN)) as taken from the Catálogo de Plantas y Líquenes de Colombia (Bernal et al., 2019). Outliers (3) have been removed.

**Table 3.** Dunn’s test results for post-hoc comparisons between categories of national vulnerability status and count of local names.

| National vulnerability status<br>category 1 | National vulnerability status<br>category 2 | Z | P unadjusted | Bonferroni P<br>adjusted |
| --- | --- | --- | --- | --- |
| EN | LC | -0.265 | 0.791 | 1.000 |
| EN | NE | 1.177 | 0.239 | 1.000 |
| LC | NE | 3.184 | 0.001 | 0.022 |
| EN | NT | 0.779 | 0.436 | 1.000 |
| LC | NT | 1.386 | 0.166 | 1.000 |
| NE | NT | -0.209 | 0.834 | 1.000 |
| EN | UN | 2.003 | 0.045 | 0.677 |
| LC | UN | 3.396 | 0.001 | 0.010 |
| NE | UN | 1.630 | 0.103 | 1.000 |
| NT | UN | 1.273 | 0.203 | 1.000 |
| EN | VU | -0.419 | 0.675 | 1.000 |
| LC | VU | -0.294 | 0.769 | 1.000 |
| NE | VU | -1.501 | 0.133 | 1.000 |
| NT | VU | -1.128 | 0.259 | 1.000 |
| UN | VU | -2.203 | 0.028 | 0.414 |

## 4. Discussion

We sought to explore the relationship between the endemic status of plants and artisan crafts in Colombia, and the relationship between the most well-known artisan plant species and the most threatened, according to international and national vulnerability statuses. Even though political boundaries do not necessarily coincide with biogeography and do not accurately represent species ranges, there is benefit to using political boundaries because conservation governance happens at political scales, as countries set budgets and design transboundary conservation initiatives (Dallimer & Strange, 2015; Echeverri et al., 2023; Gentili, 2011). Moreover, cultural exchange is dynamic, moving in, across, and despite borders, as is the exchange of biological species (Dong et al., 2017; Crosby, 2003; Mack & Lonsdale, 2001).

### 4a. Craft, Food, Medicine, and Plant-Human History

Artisan crafts are a major component of all human cultures, as is food and medicine, all three of which are heavily reliant on plant species (Balick & Cox, 1996), and can provide context to the spread and decline of culturally important plant species. Basketry is an artisan skill that has been practiced in the American continents for thousands of years by multiple Indigenous peoples, for practical, aesthetic, political, and spiritual uses (Geib & Jolie, 2008; Anderson, 1996; Guss, 1989; Balick & Cox, 1996). In Colombia, the chumba wumba palm (*Astrocaryum standleyanum*) and patawa (*Oenocarpus batua*) are both regional palms with native distributions from Panama into the Amazon; both are used for basket weaving (García et al., 2013; Smith, 2015). Further, the chumba wumba palm has many documented uses in artisan crafts, beside basketry (i.e. hats, hammocks, mats), and has been adopted by non-Indigenous populations as well, demonstrating how the cultural importance of the palm has expanded to multiple peoples as a result of the socio-political history of the region (Fadiman, 2008; García et al., 2013; Potvin et al., 2003; Runk, 2001). While native plant species may hold special cultural importance in a culture, introduced species may become favored over native species for various reasons (Bennett & Prance, 2000; Cadena-González, Sørensen, & Theilade, 2013; Hart et al., 2017). The adoption of introduced plant species into medicinal practice is partly due to the introduction of non-native food species, that also had medicinal use (Bennett & Prance, 2000). It is widely known that food species have been cultivated and spread by human action, like corn, a crop that was domesticated approximately 7,000 to 10,000 years ago from teosinte (*Z. mexicana* and *Z. perennis*) in the highlands of current day Mexico; its variations quickly spread across the two continents (García-Lara & Serna-Saldivar, 2019; Yang et al., 2023). Corn was also one of the three outliers for number of local common names attributed to each species.

The three outliers—the calabash tree (*Crescentia cujete*), the bottle gourd (*Lagenaria siceraria*), and corn (*Zea mays*)—are all well-known, widespread plants in Colombia and internationally. This is shown in the number of local common names attributed to each species (45, 50, and 52, respectively) and the number of departamentos listed alongside those names (29, 26, and 30 departamentos out of a possible 32, respectively). The calabash tree is native to Colombia and a regional species. Corn was introduced thousands of years ago and so we may consider it made regional by human action. Likewise, the earliest bottle gourd in the Americas dates back to approximately 10,000 years ago before present (BP) according to mass spectrometer radiocarbon dating, despite being native to Africa (Lim, 2012). While these three outliers point to ancient human interactions, the presence of some introduced species in Colombian artisan crafts (i.e. *Abrus precatorius*, *Chrysopogon zizanioides*, *Coix lacryma-jobi*, *Fraxinus chinensis*, and *Cocos nucifera* to name a few) may be the result of more recent global events, including but not limited to Spanish settler-colonialist campaigns, the globalization of trade in the last two to four hundred years, and accidental introduction by individual travel (Crosby, 2003; Bennett & Prance, 2000; Mack & Lonsdale, 2001).

This is to say, again, that culture is dynamic; the result of many ever-changing, intertwined history and relationships between peoples and the lands they rely on. While we may see regional and introduced species more represented in documented artisan crafts than native species, these results must be considered within the context of Colombia and the nation’s history. Ultimately, it would be inappropriate to theorize without evidence of the specific intricacies of human-plant relationships.

### 4b. Endemic Artisan Plant Species

While plant species endemic to Colombia are present in artisan crafts, artisan crafts in Colombia are not heavily reliant on endemic plant species. Instead, plant species found across the northern South American and Central American region were heavily represented. <u>As were</u> introduced species, though less so than regional ones. This might be partially explained by small population size and rarity. Many introduced plant species are effective at pushing out native species, and so exacerbate endemic rarity and threaten traditional craft practices (Ticktin, Whiteheadm, & Fraiola, 2006; Pfeiffer & Ortiz, 2007; Pfeiffer & Voeks, 2008). The small proportion of endemic plant species in documented artisan crafts may not be representative of historical proportions, or of preference of use. Further, the cultural awareness of endemic plant species may be greater than what can be shown with documented production (Reyes-García, 2006).

### 4c. Vulnerability Status and Cultural Awareness

There is a significant relationship between a species’ national vulnerability status and the number of associated local common names (Figure 3). With the number of local names as a proxy for cultural awareness and potential importance, this result suggests that, in general, locally well-known plant species are more likely to be given a higher vulnerability status (VU and EN) and the least locally known plants are more likely not to be evaluated (NE) or not even profiled on the national vulnerability list as an existing plant in need of evaluation (UN). Like with endemic plant species, we at first inferred the small number of local common names for UN categorized plants may indicate that these plant species are rare, and so easily missed in national vulnerability assessments. However, a closer look at this category revealed that only six plant species were not profiled and of these six, three were introduced to South America (*Agave sisalana*, *Fraxinus chinensis*, *Salix viminalis*), two were regional (*Pontederia crassipes*, *Astrocaryum tucuma*), and one was introduced to Colombia but regional to South and Central America (*Tillandsia utriculate*). Instead of rarity and lack of awareness for the species, the missing profiles for these plants may be because of a choice to exclude introduced plants, or simple oversight.

Even excluding UN species, we still see differences across groups between number of local common names and national vulnerability status for all species given a status (NE to EN). The similar range in the number of local names for NE and LC, and similar range size for NT, VU, and EN, suggest some additional factor deciding how plant species are being categorized at the national level. We suspect that many plant species important to Colombian artisan crafts have not been thoroughly studied for recent population changes in size or health. Like IUCN assessments, national assessments may be opportunistic with data sourcing and, as a result, sparsely researched plants may not be assessed or re-assessed due to a lack of information.

One possible alternative explanation for the positive relationship between number of local common names and national vulnerability status may be that most of the documented plants are regional, and so may be so widespread that the species is considered of little conservation concern or not yet in need of evaluation. Lopez-Gallego & Morales-Morales (2023) found that endemic and threatened species of trees were not uniformly distributed across the biographical regions of Colombia, and for trees in the Amazonia and Guyana shield regions, most species were labeled as Least Concern by the IUCN, because of large extant of occurrence (EOO)—the estimated minimum convex polygon that encompasses all known occurrences of a species—and lack of threats (Lopez-Gallego & Morales-Morales, 2023). The plant species included in this study may also have been assessed as Least Concern for these reasons. But, while a culturally important plant species may not be threatened on a global or continental scale, if that species is lost locally then local people’s ability to continue practicing their craft may be threatened.

### 4d. Limitations

Throughout the data collection process, we thought critically about what information we were gathering from our sources, considering such things as from where and when the information was coming, potential biases in sources, source goals and assumed audience, languages biases (i.e. Spanish and English represented more than, or preferred over Indigenous languages), and geographic biases (i.e. overrepresentation of more populous departments). For instance, Artesanías de Colombia, Universidad Nacional de Colombia (UNC): Catálogo de Plantas y Líquenes, and UNC: Nombres Comunes were all primarily in Spanish, while the Catalogue of Useful Plants of Colombia, Plants of the World (POWO), and the IUCN Red List were all in English. Data collection was done by bilingual Spanish and English authors, with both languages used throughout the process. Many of the artisan crafts documented in Artesanías de Colombia are Indigenous, and the underrepresentation of Indigenous languages in the accessed archives may have affected this study in several ways. For example, some endemic artisan plant species may have names in Indigenous languages only, and the underrepresentation of these languages may have led to the unintentional exclusion these species from our study. Some of the included species may also, in reality, have more associated common names than recorded.

This study is also limited by the archival methods employed. No in-person research was conducted, nor did we collaborate with or work alongside all the cultural communities who are included in this research by way of their craft.

We also acknowledge the discussion regarding the distinction between “art” and “craft”, and how the distinction of the two has come, at least in part, from a history of elitism, classism, racism, sexism, and other social influences (Markowitz, 1994). For the purpose of this paper, we use the terms artisan crafts because it is under these terms that data was organized by our sources.

### 4e. Conservation Implications and Future Research

This study was inspired by calls to conserve ecosystems according to biocultural values—protecting biodiversity for the relationships between people and the natural world as much as for ecosystem function, evolutionary history, and biodiversity for biodiversity’s sake (Bond et al., 2019; Gavin et al., 2015; Rozzi, 2013; Reyes-García et al., 2023). However, we acknowledge the present critiques for bioculturalism as a term and conservation goal (Rashkow, 2014). Rashkow (2014) makes the argument that biocultural conservation can still fall back on colonial ideas and stereotypes that paternalistically call to “protect” Indigenous peoples, as if they do not have their own voices and must be spoken for, or worse, are akin to animal and plant species in need of “preservation” (Rashkow, 2014). We acknowledge Rashkow’s (2014) critiques and call to interact with the history of conservation in an honest and critical manner. With this critical perspective and the results of this study, we encourage future studies—especially in conservation research—to cultivate collaborations that represent multiple fields of knowledge, including ecologists, anthropologists, local and Indigenous community members, and practiced experts in craft and other cultural practices (Singeo & Ferguson, 2023). Such collaboration should prioritize the goals and concerns of the communities involved (Polfus et al., 2016; Reid et al., 2021). Extra effort on behalf of researchers should be taken to understand the traditional knowledge and values systems of a collaborating community (Polfus et al., 2016; Reid et al., 2021; Shackeroff & Campbell, 2007). Conservation practitioners should continue to incorporate cultural systems and practices as a part of deciding conservation priorities and policy.

## Conclusion

Human beings are animals of our environment, just like any other. Our cultures developed alongside the many plant and animal species now threatened with extinction. Biodiversity loss is both a crisis of ecological function as it is a crisis of identity—who are we when the lands and waters that homed us are no longer there? This is not a novel question, as there are many who have approached from the perspective of settler-colonialism, imperialism, diaspora, and the like. Ultimately, biodiversity loss is one part of a larger environmental crisis that affects people unequally across the globe. If we do not confront and account for this inequity at every stage of conservation, then we are complacent in it. Ecological metrics are vital to prioritizing species for conservation, we do not dispute this. However, there is no ecological metric capable of showing which species will be more deeply mourned. There is no ecological metric that can measure the weight of absence. The results of this study show that many plant species important for Colombian artisan crafts are not being assessed for vulnerability to extinction, and even the most well-known artisan plant species lack profiles on the IUCN Red List international conservation. A species can be given a profile on the IUCN Red List before a complete assessment; the lack of a profile on such a highly utilized, international conservation resource may render a species invisible to international aid. We, the authors, hope this study may encourage culturally informed conservation, and serve as a reminder of all the parts of us that may still be lost.

## Acknowledgements

We acknowledge that we wrote this manuscript from the unceded territories of the Gabrielino Tongva and the Ohlone peoples. We celebrate the perseverance of all Indigenous Peoples across the world who stand strong in preserving their cultural traditions, such as the ones we write about in this manuscript. We are grateful for conversations with the Blumenstein Lab, and Tenzin Norzin, Stacie Wolny, Dr. Hector Angarita, Dr. Tong Wu, and profs. Gretchen Daily, Tadashi Fukami, and Rodolfo Dirzo who helped shape this project since its inception. We also thank Colin O’Brien-Lux for sparking the connections between K.V.H. and A.E..

## APPENDIX. SUPPLEMENTARY DATA

**Table S.1.** Data sources consulted for this project.

| Data source | Information in the source | Search terms used | Reference |
| --- | --- | --- | --- |
| Artesanas de Colombia | Prime plant & animal material used in making documented artisan crafts; kinds of artisan crafts (i.e. basketry, wood working, natural dyes); common and scientific names of prime material; additional context for plant identification. | Prime Materials (Materias Primas) | Centro de Investigación y Documentación para la Artesanía – CENDAR Artesanías de Colombia (n.d.). CENDAR – Biblioteca Digital. Artesanías De Colombia. https://cendar.artesantiasdecolombia.com.co/ |
| Universidad Nacional de Colombia – Catálogo: Nombres comunes | The largest collection of Colombian plant common names ever documented. Common names local to the nation of Colombia were extracted from this archive and found by searching by scientific name at the species level. | Common names (Nombres comunes) | Bernal, R., Galeano, G., Rodriguez, A., Sarmiento, H., & Gutierrez, M. (2017). *Nombres Comunes de las Plantas de Colombia*. Universidad Nacional de Colombia. http://www.biovirtual.unal.edu.co/nombres-comunes/en/ |
| Universidad Nacional de Colombia - *Catálogo de Plantas y Líquenes de Colombia* | A digital archive of 28,000 plant and lichen species found in Colombia. Archive was searched by scientific name, at the species level. Information on national vulnerability status for each profiled plant species was extracted from this archive. | Search by scientific name. | Bernal, R., S.R. Gradstein & M. Celis (eds.). 2019. *Catálogo de plantas y líquenes de Colombia*. Instituto de Ciencias Naturales, Universidad Nacional de Colombia, Bogotá. |
| Catalogue of useful plants of Colombia | This book was made in partnership with various Colombian and European contributors to study and catalog useful plants of Colombia. There are twelve chapters of research included, and a catalog of 7,472 plants. Each catalog entry contains available information on supraspecific taxa, species, common names, species origin, geographic information, conservation status, and use. All of which was used for this study where applicable, and, or, to verify plant identification at the species level. | Endemic status | Diazgranados, M., Cossu, T., Kor, L., Gori, B., Torres, G., Carretero, J., ... & Negrão, R. (2022). Annotated checklist of useful plants of Colombia. In *Catalogue of useful plants of Colombia*. Royal Botanic Gardens, Kew. |
| IUCN Red List | An international collaboration and openly available comprehensive list of the world's threatened animal, fungus, and plant species. Included species are assigned a global extinction risk status; possible categories are "Not Evaluated," "Data Deficient," "Least Concern," "Near Threatened," "Vulnerable," |  | IUCN (2023). The IUCN Red List of Threatened Species. Version 2023-1. https://www.iucnredlist.org/en |
|  | “Endangered,”<br>“Critically Endangered,”<br>“Extinct in the Wild,”<br>and “Extinct.” |  |  |
| Royal Botanical Gardens KEW - Plants of the World Online (POWO) | An international collaborative program and digital archive of the world’s flora, gathered over the last 250 years. Information on plant endemic, regional, and introduced status was extracted from these profiles. Information on use, scientific name at the species level, and international vulnerability status was verified using these profiles. |  | POWO (2023). Plants of the World Online. Facilitated by the Royal Botanical Gardens Kew. Published on the Internet;<br>http://www.plantsoftheworldonline.org/.<br>Retrieved 18 July 2023. |

**Table S.2.** Results of Fisher’s exact tests.

| Variable 1 | Variable 2 | Data | Fisher's Exact Test P value |
| --- | --- | --- | --- |
| Endemic status | IUCN status | All data (simulated p value, n=2000 simulations) | 0.685 |
| Endemic status | IUCN status | Excluding endemics (n=2000 simulations for p value) | 0.859 |
| Endemic status | National vulnerability status | All data (n=2000 simulations for p value) | 0.086 |
| Endemic status | National vulnerability status | Excluding endemics (n=2000 simulations for p value) | 0.142 |

**Table S.3.**
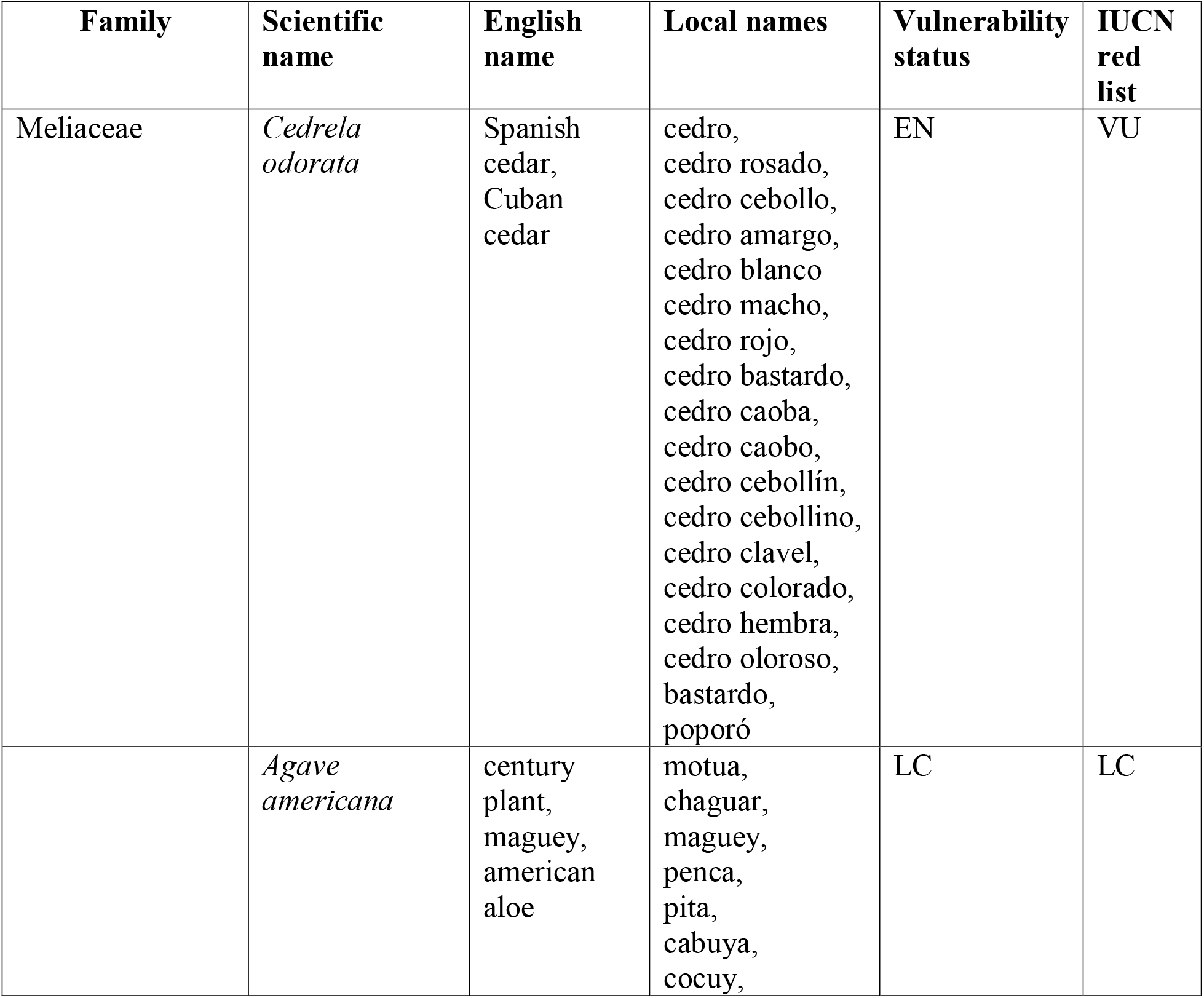
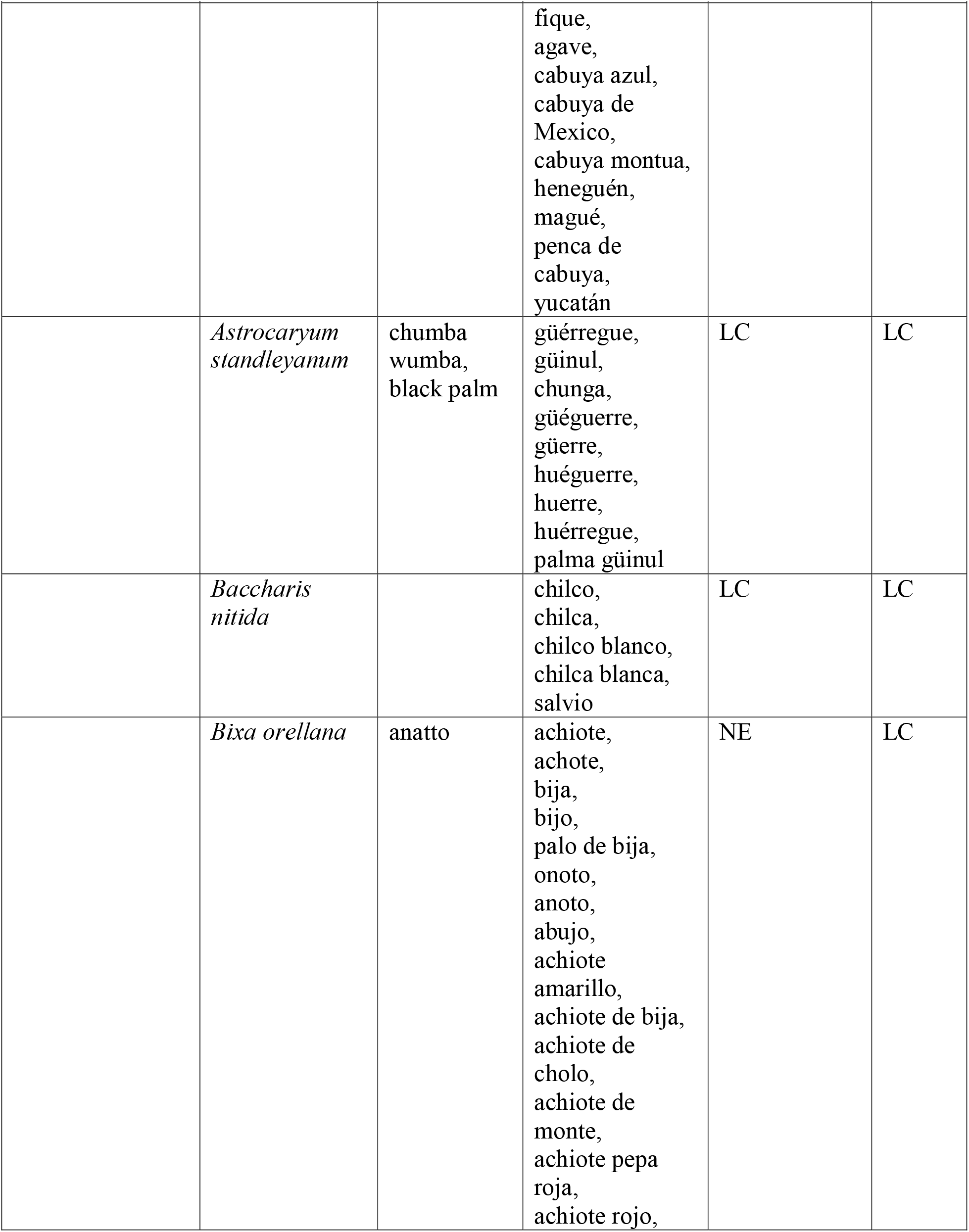
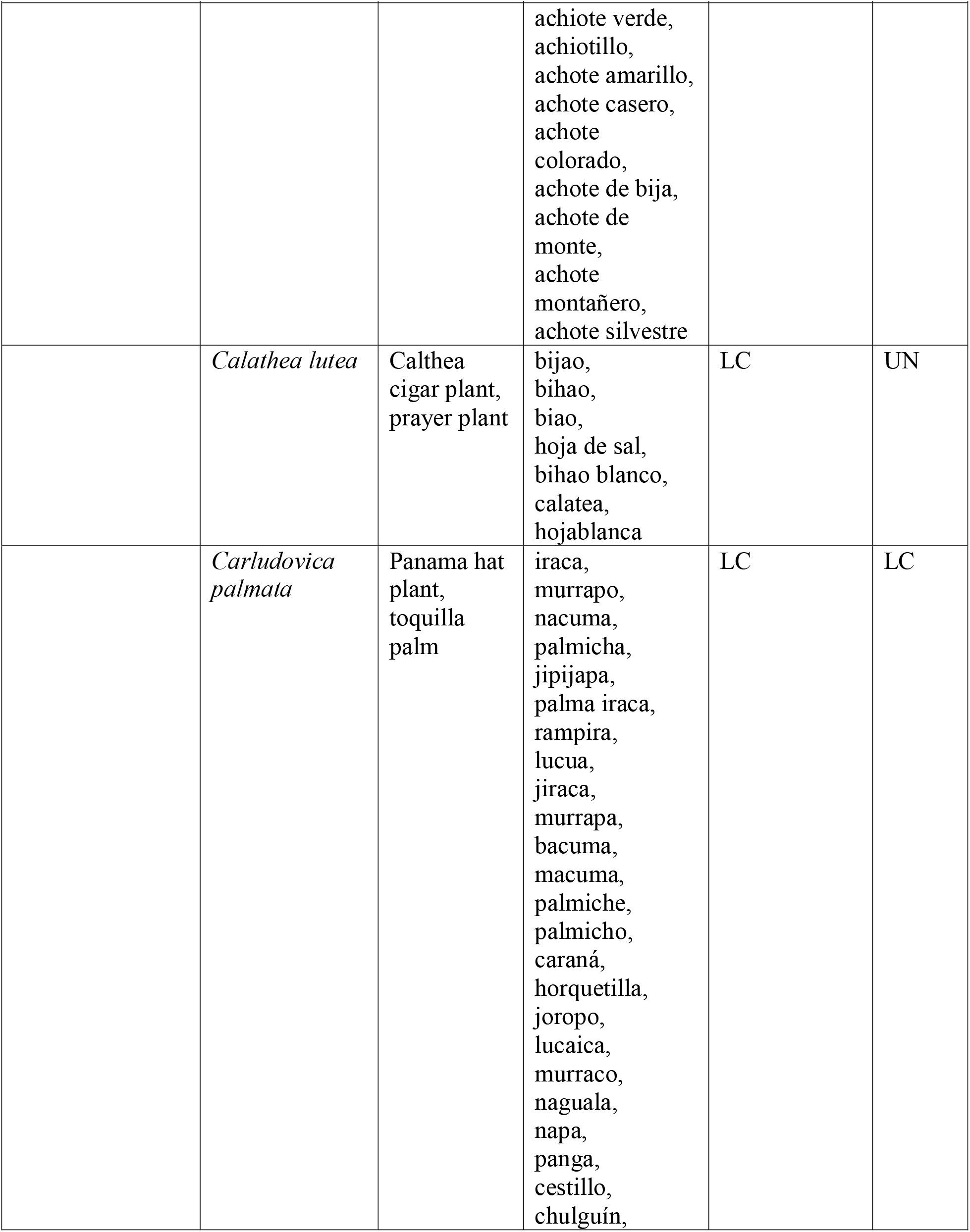
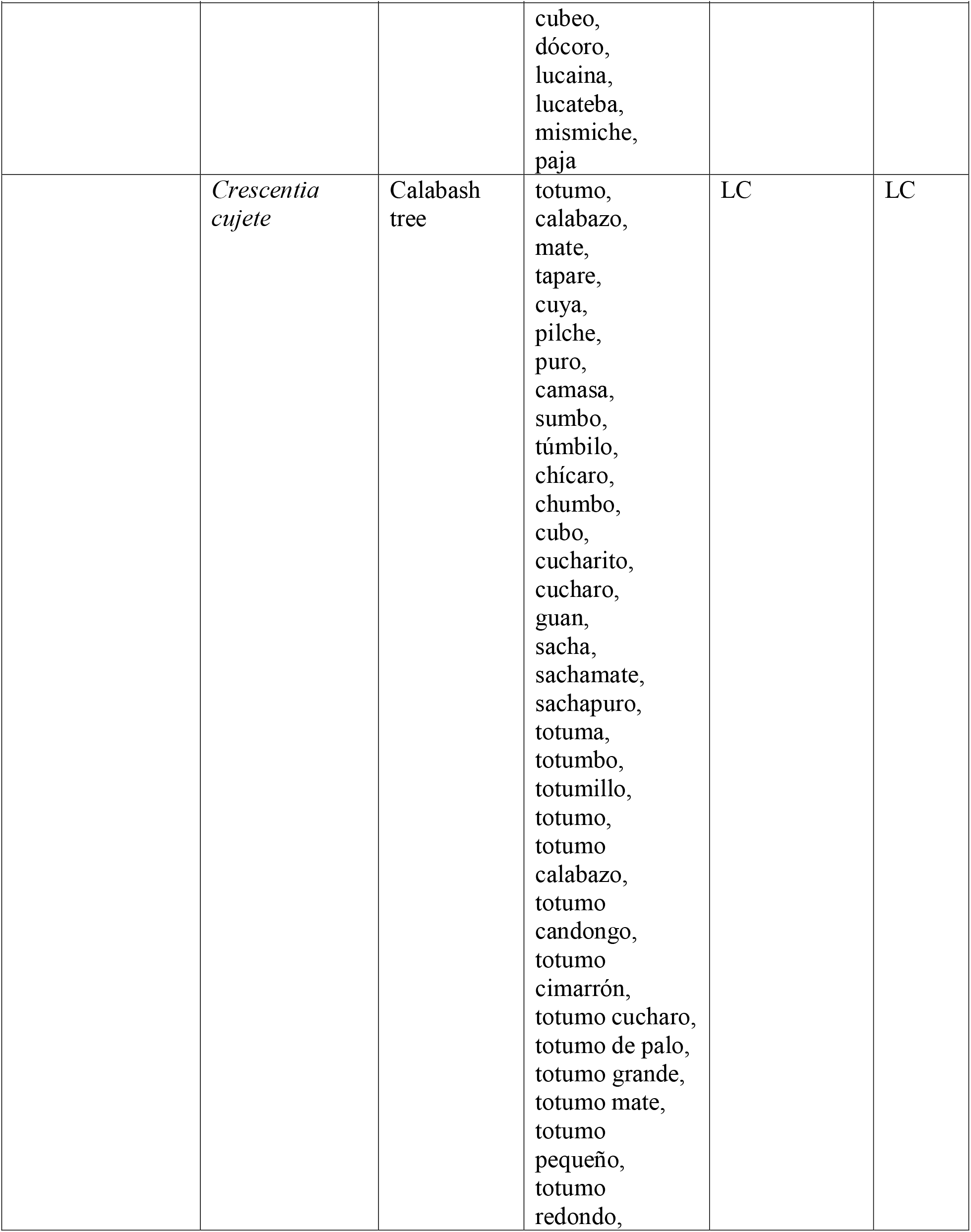
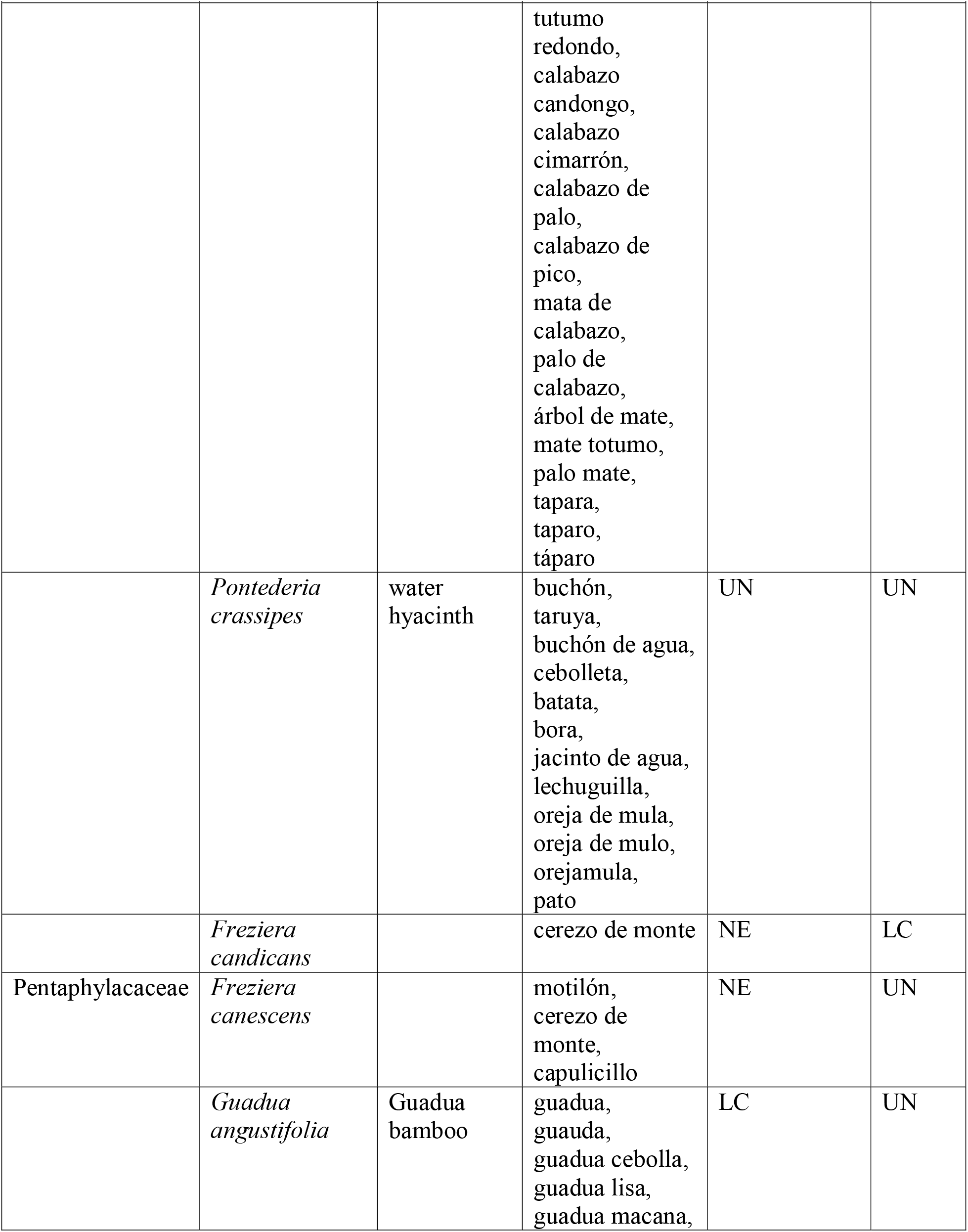
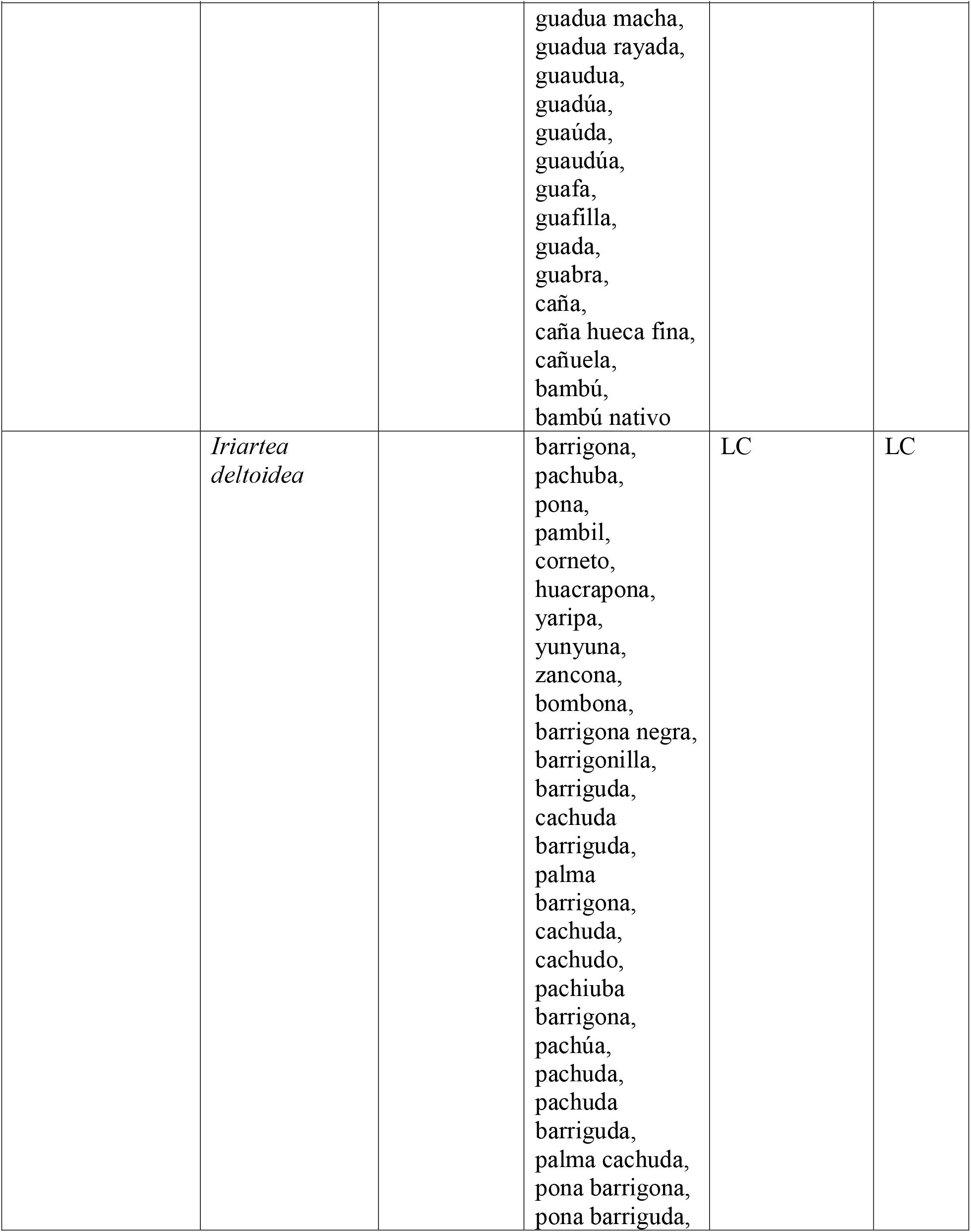
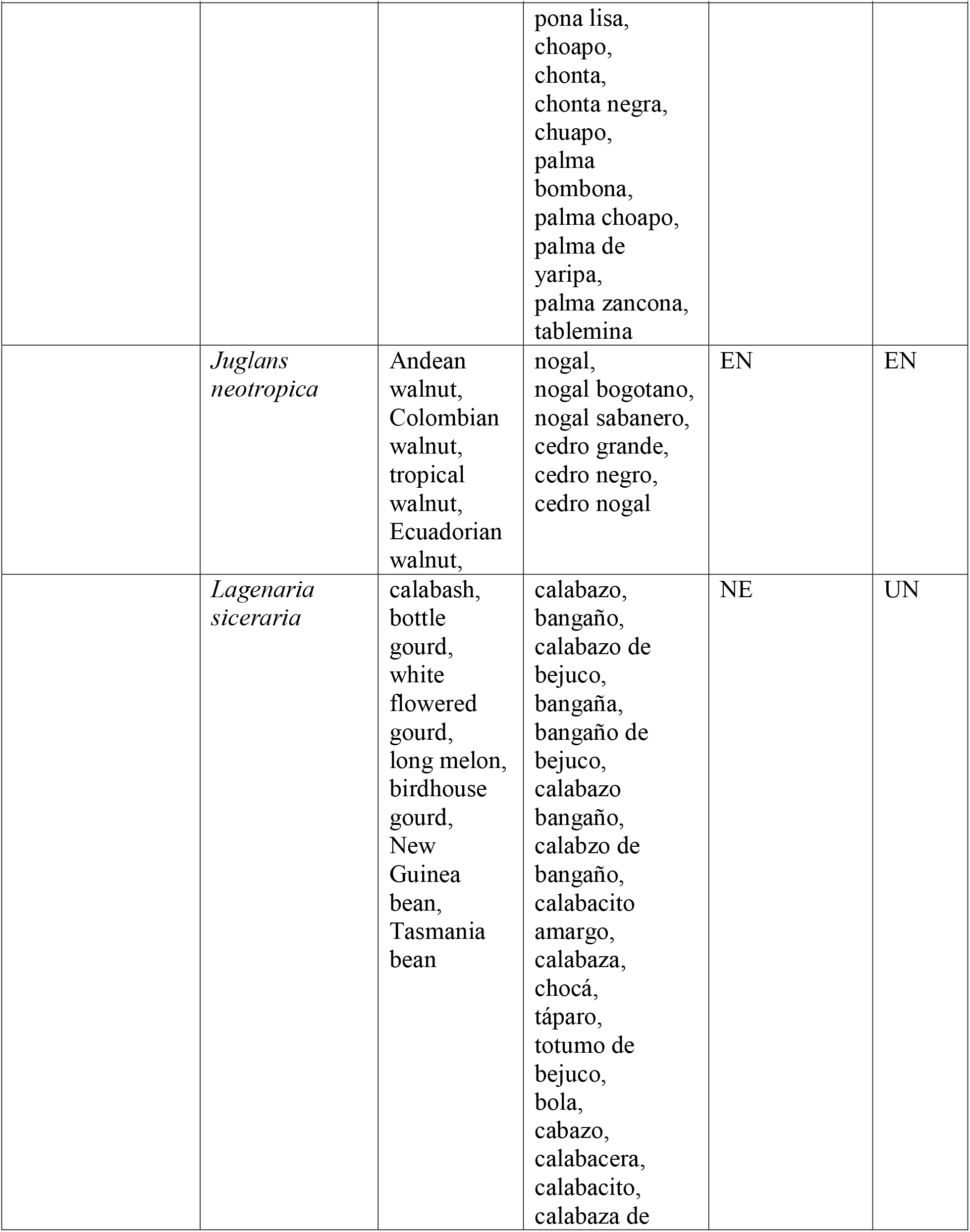
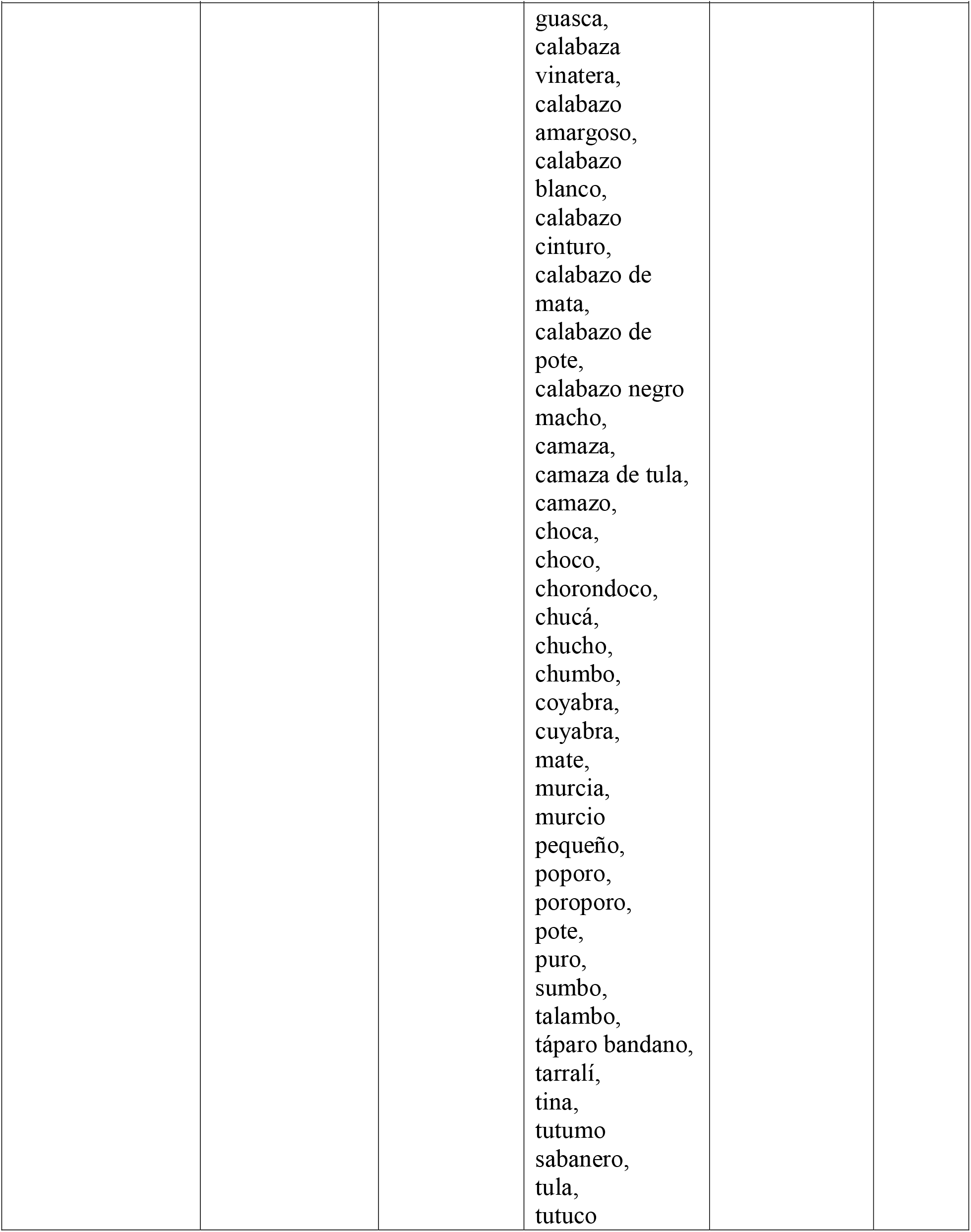
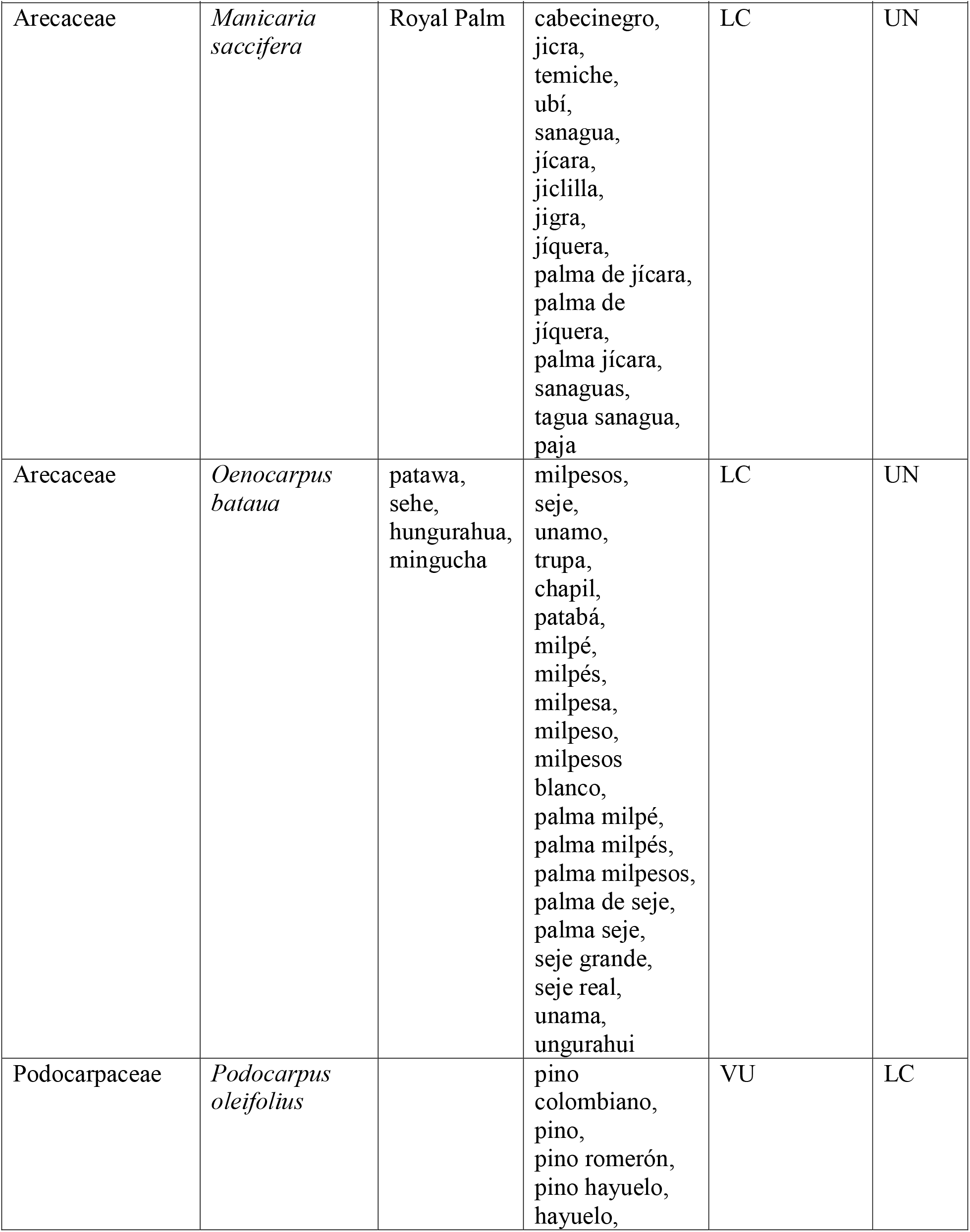
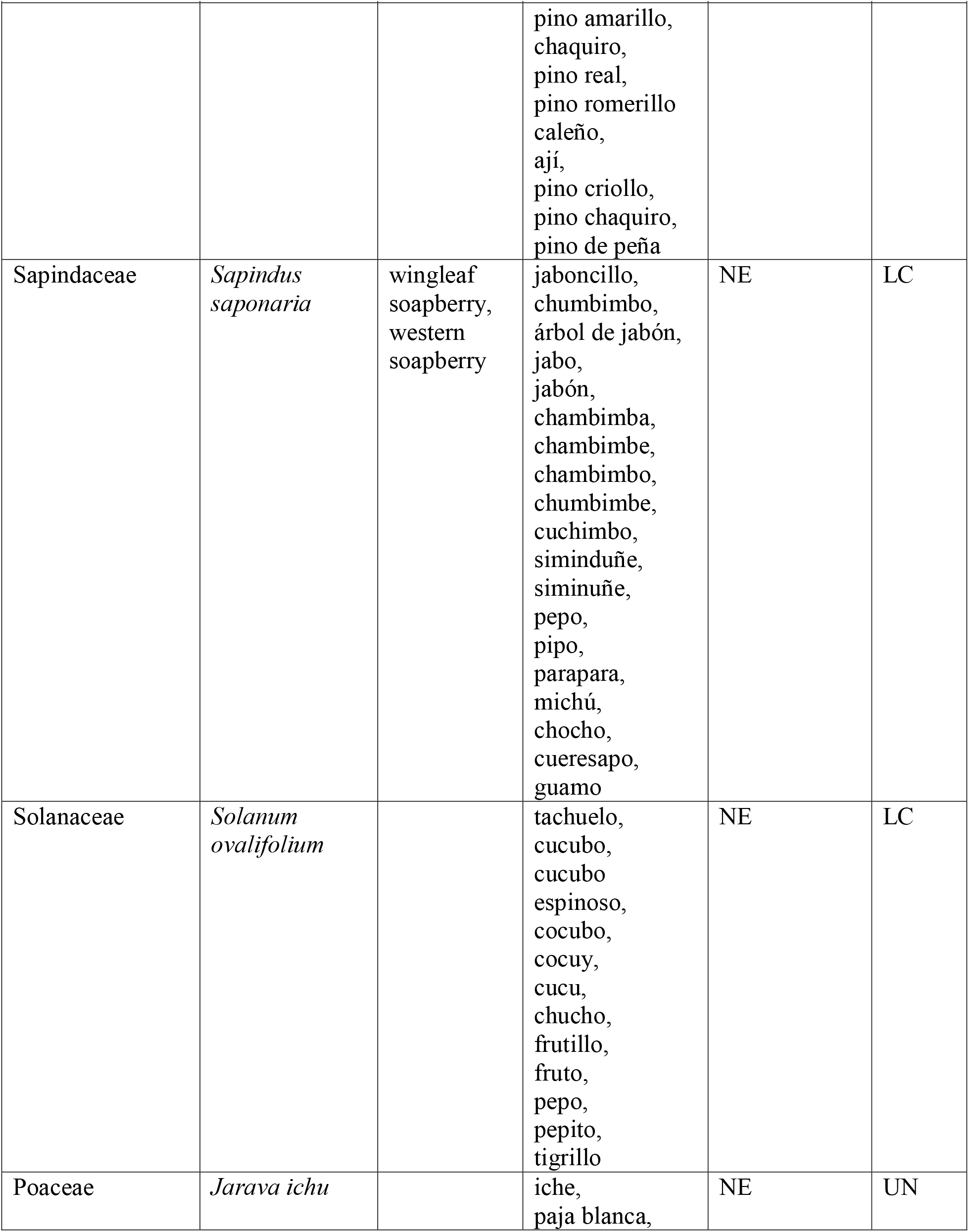
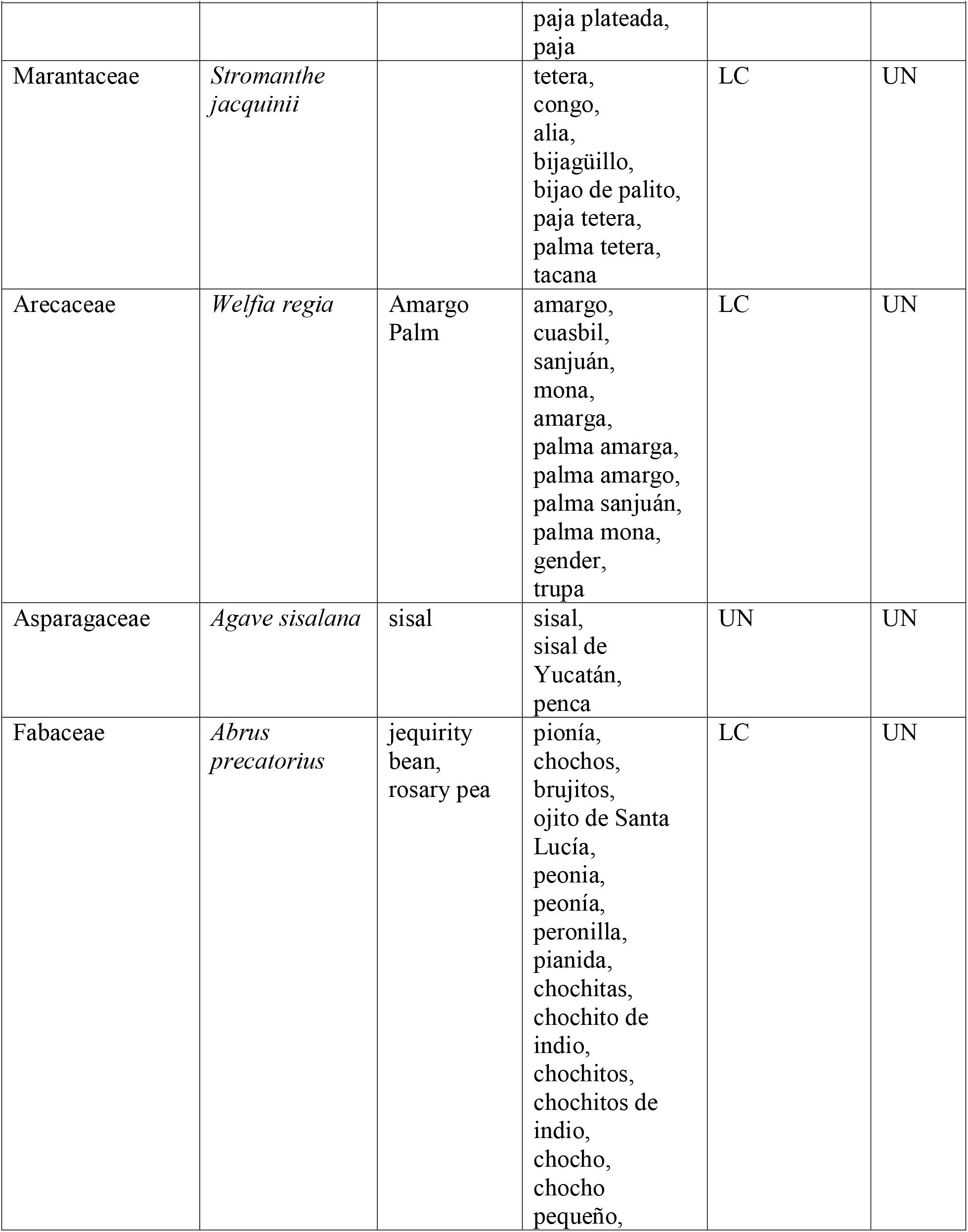
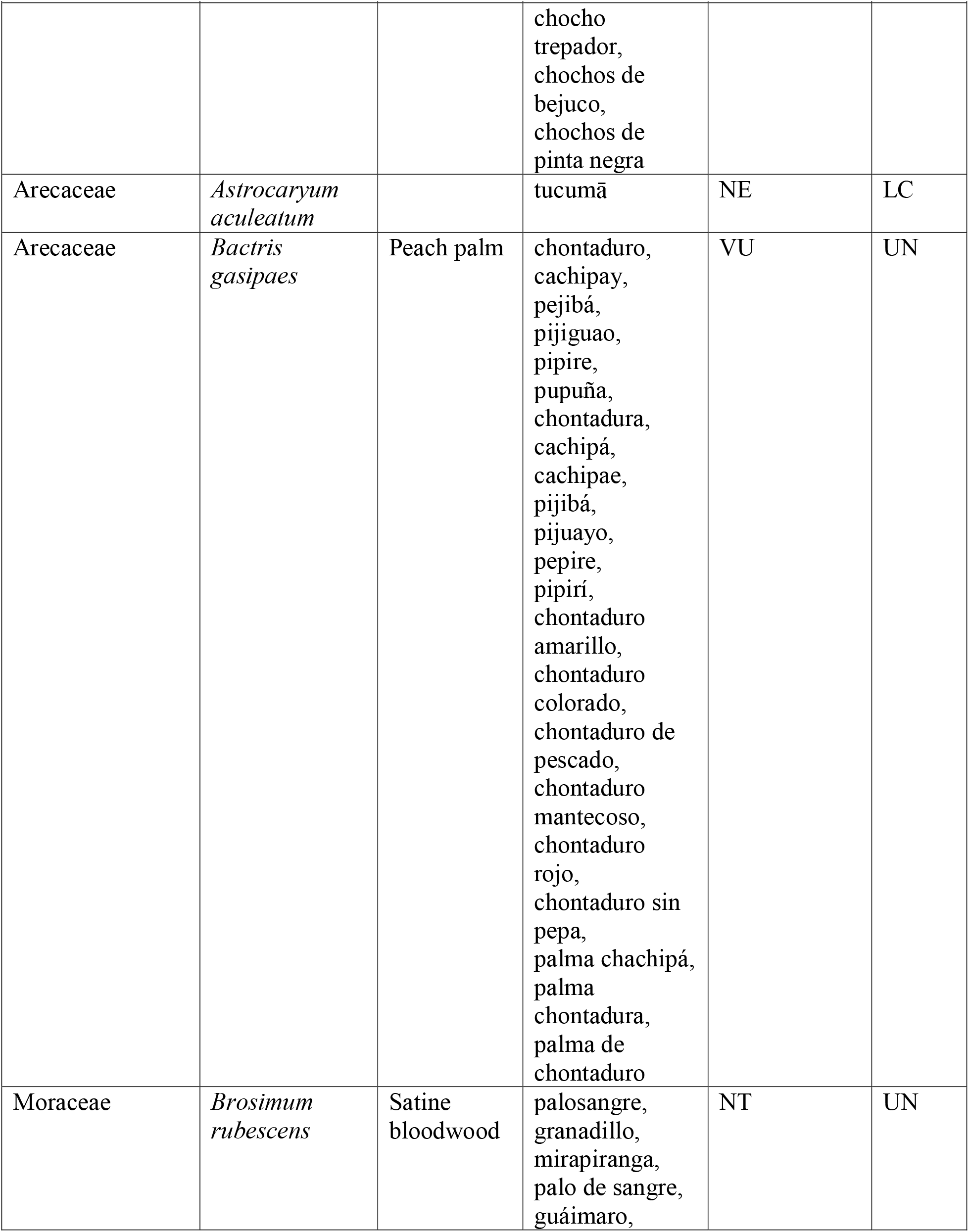
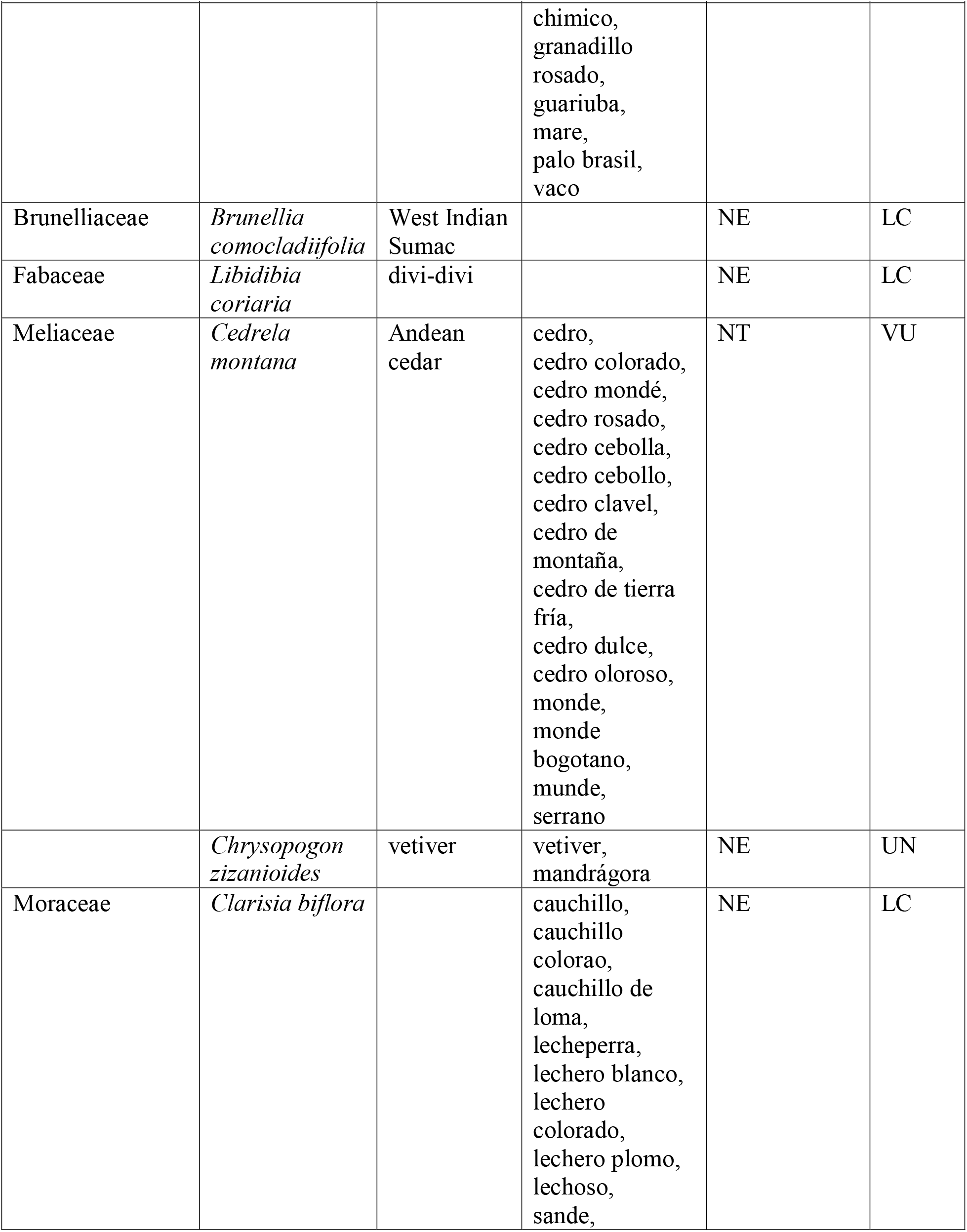
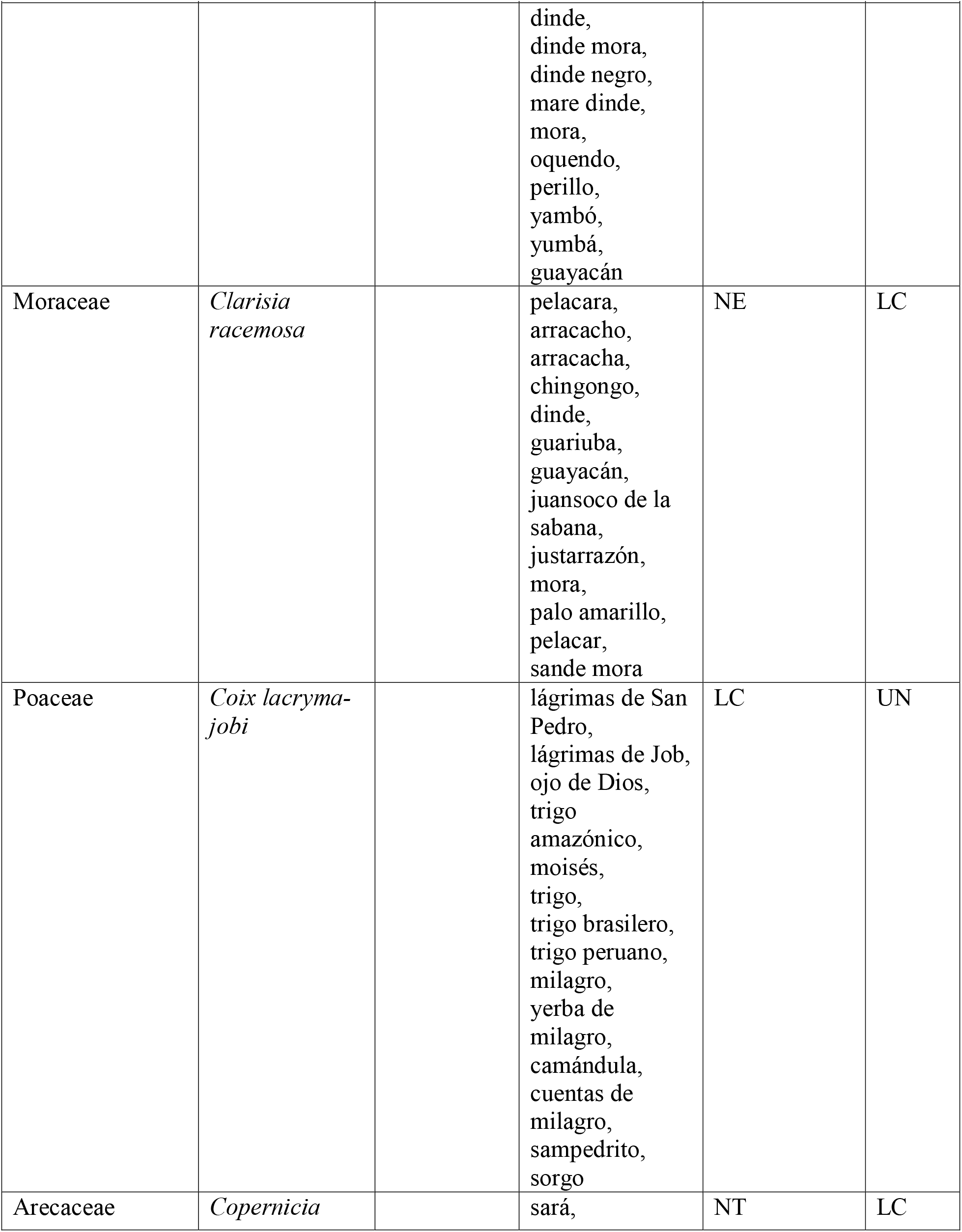
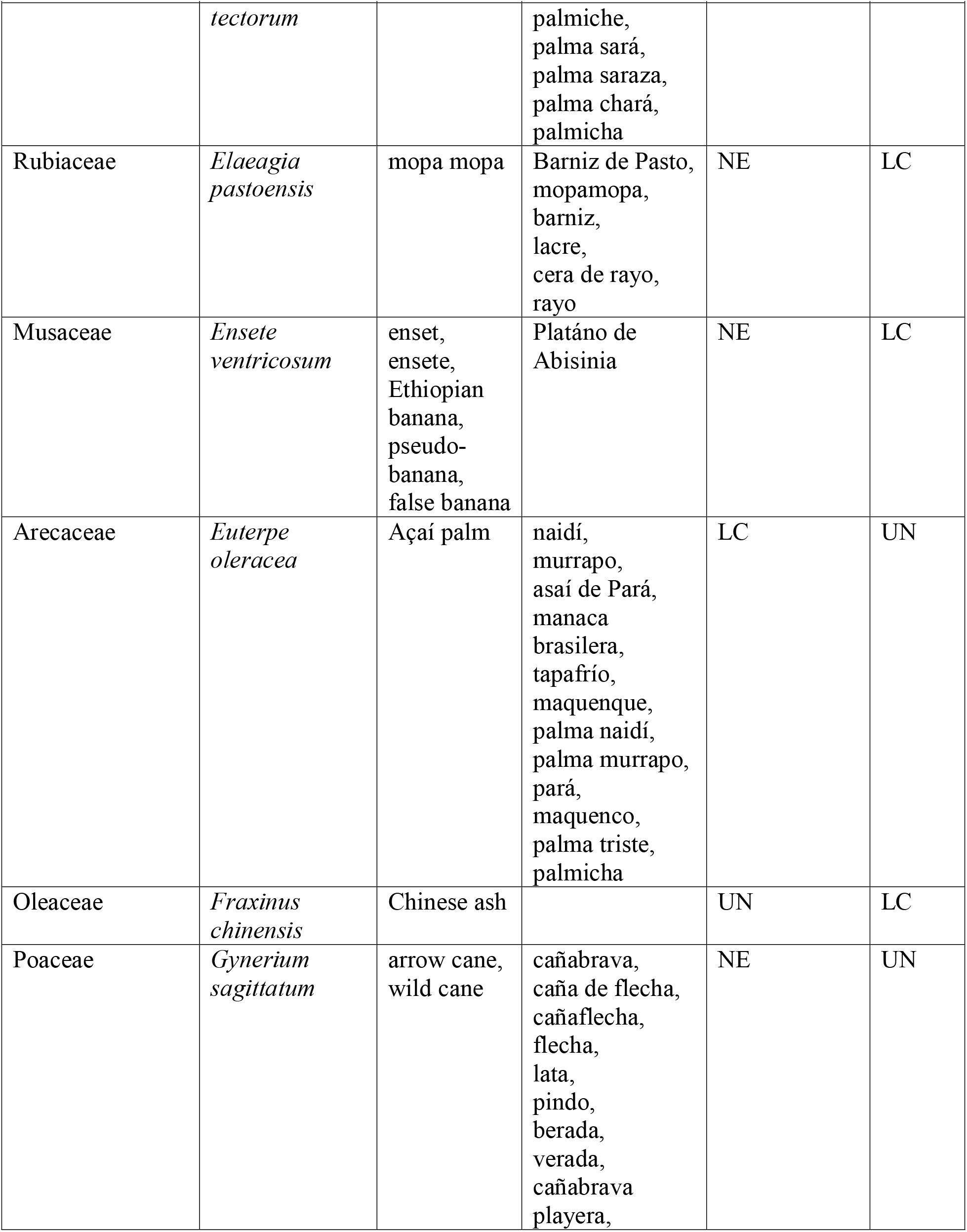
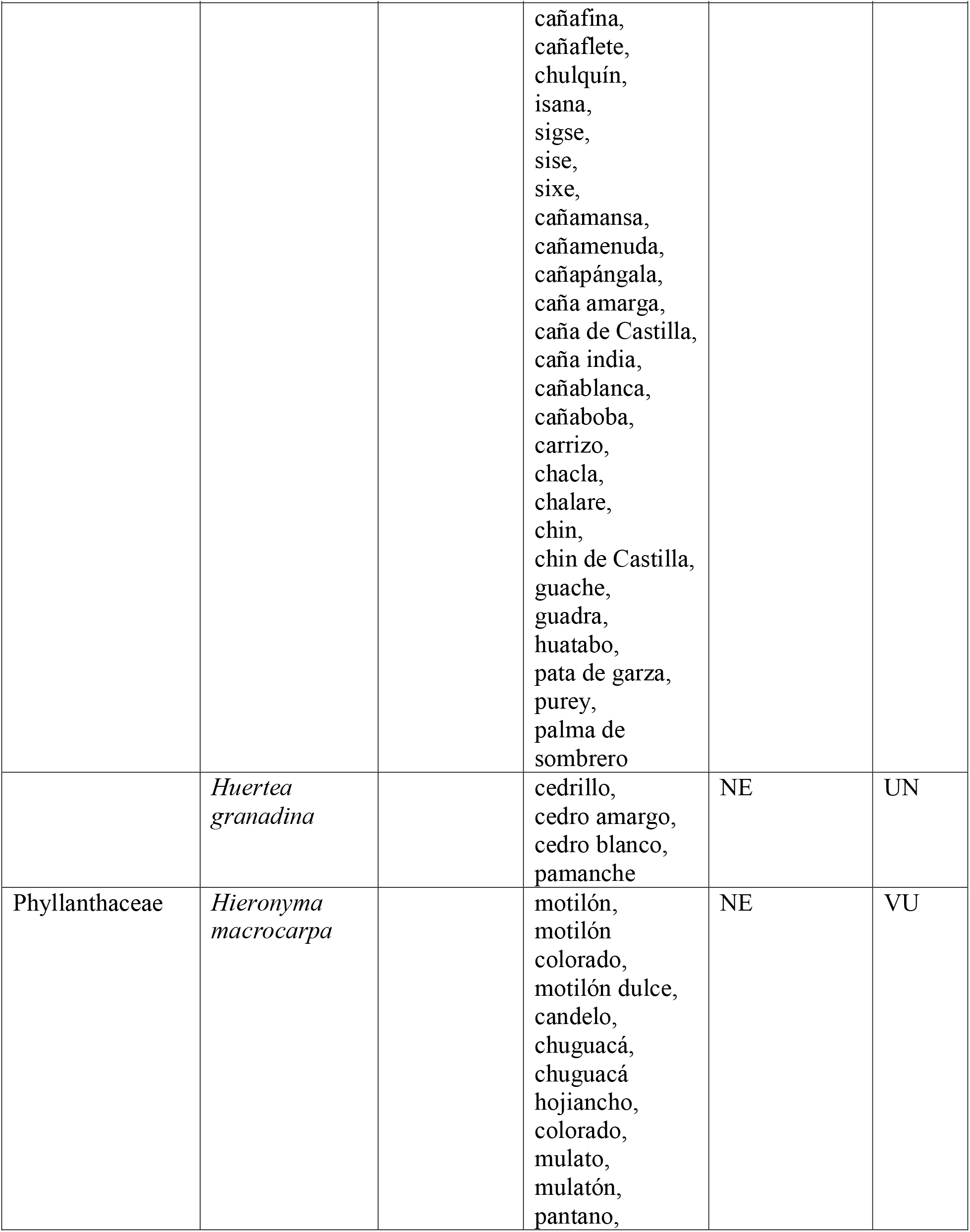
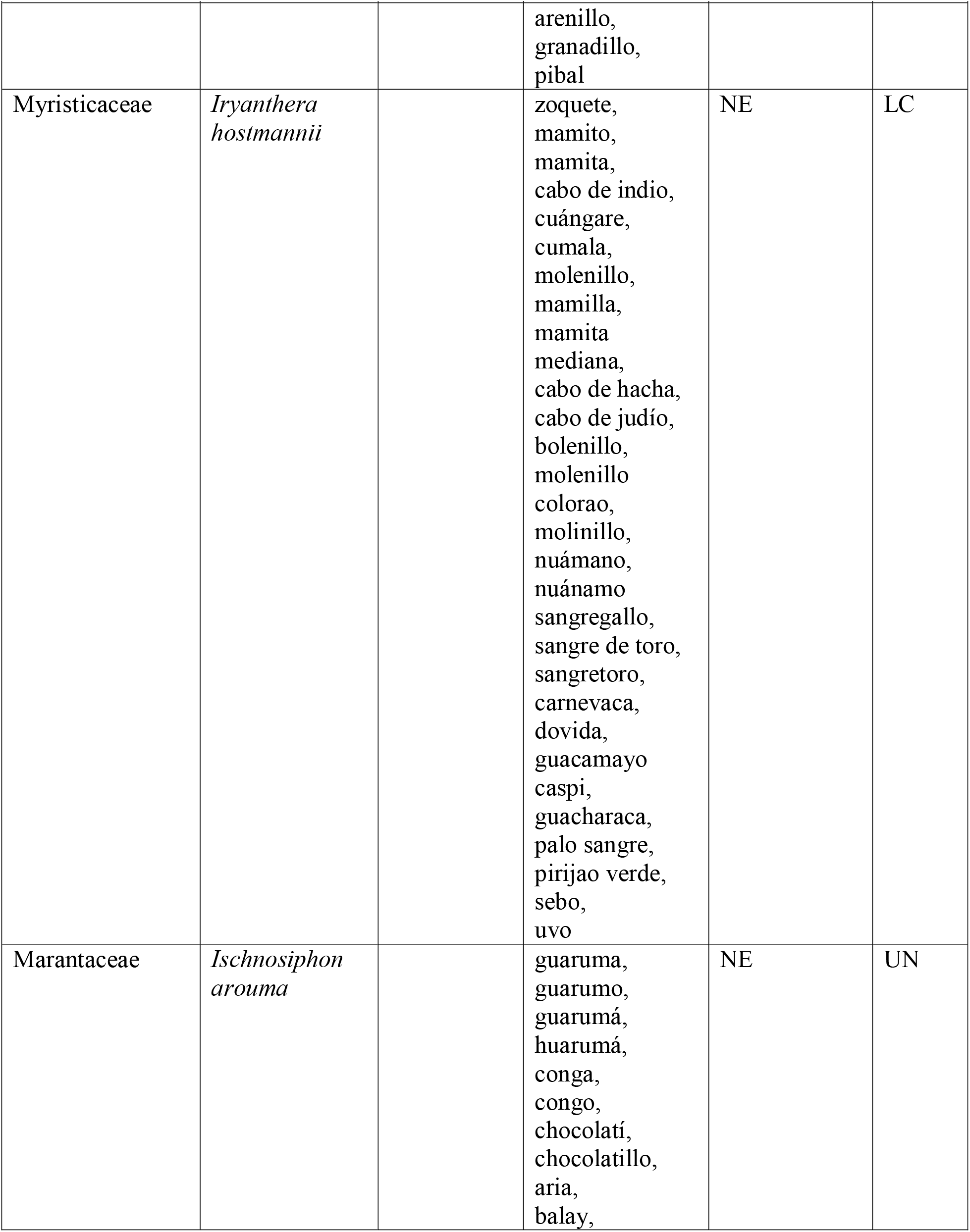
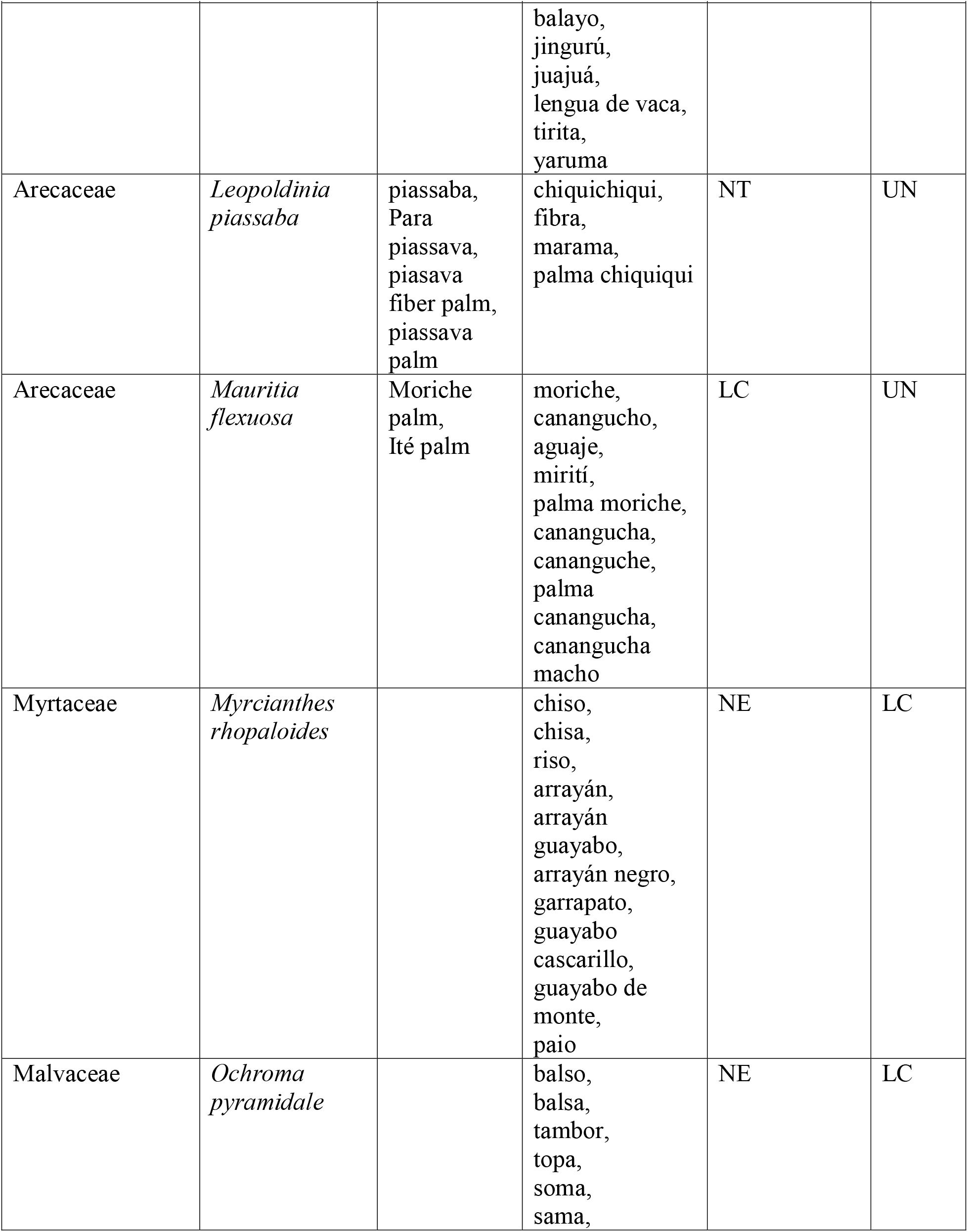
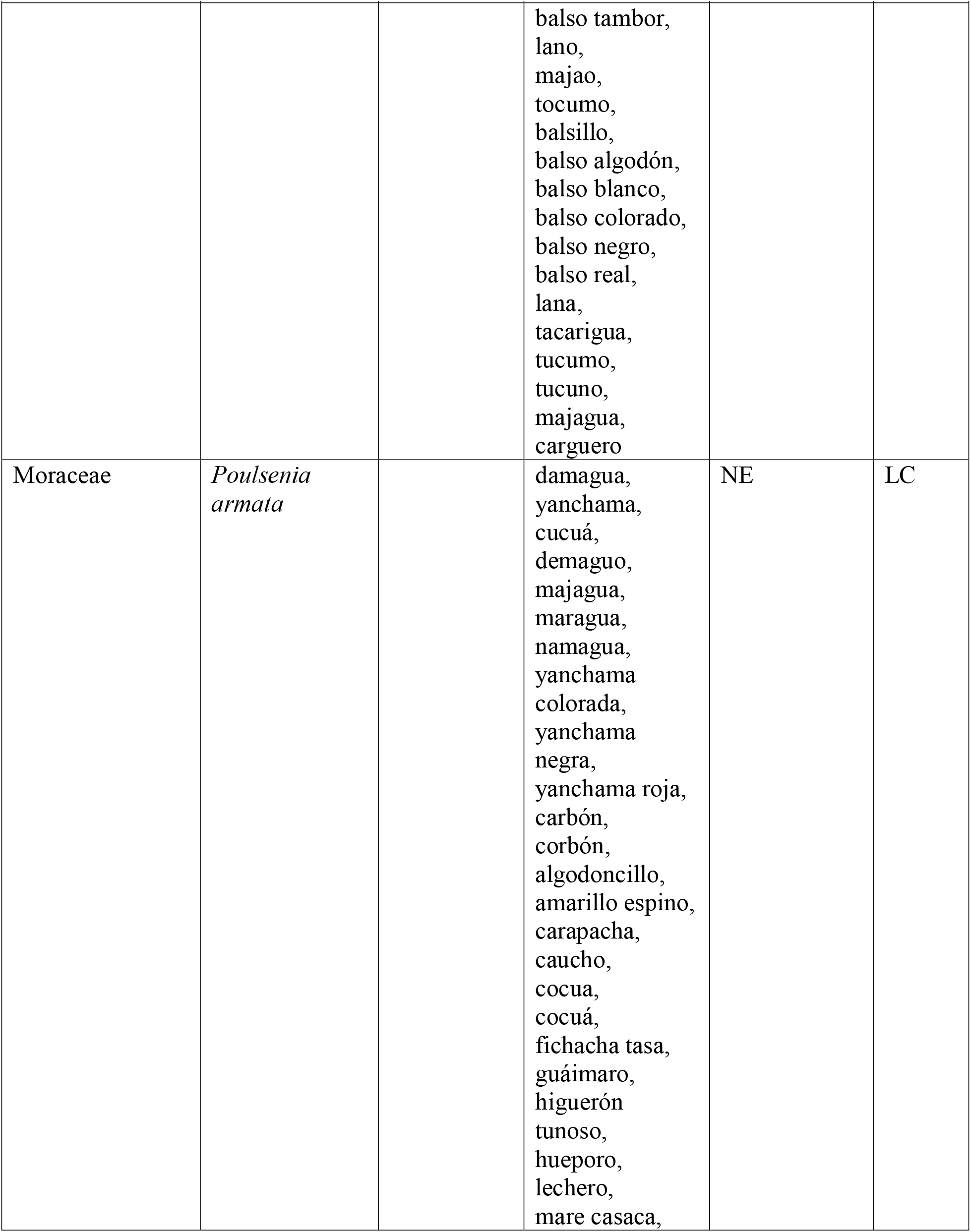
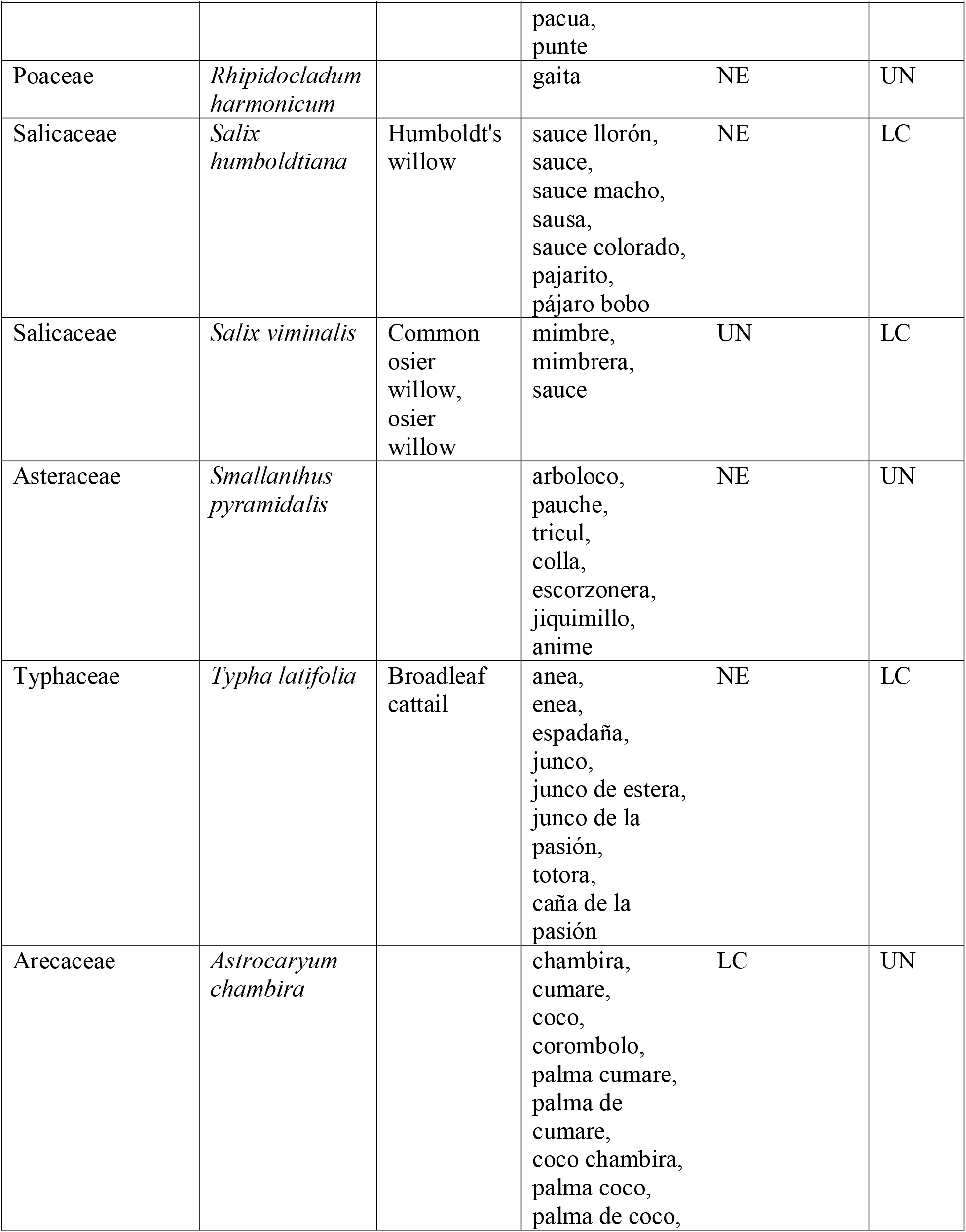
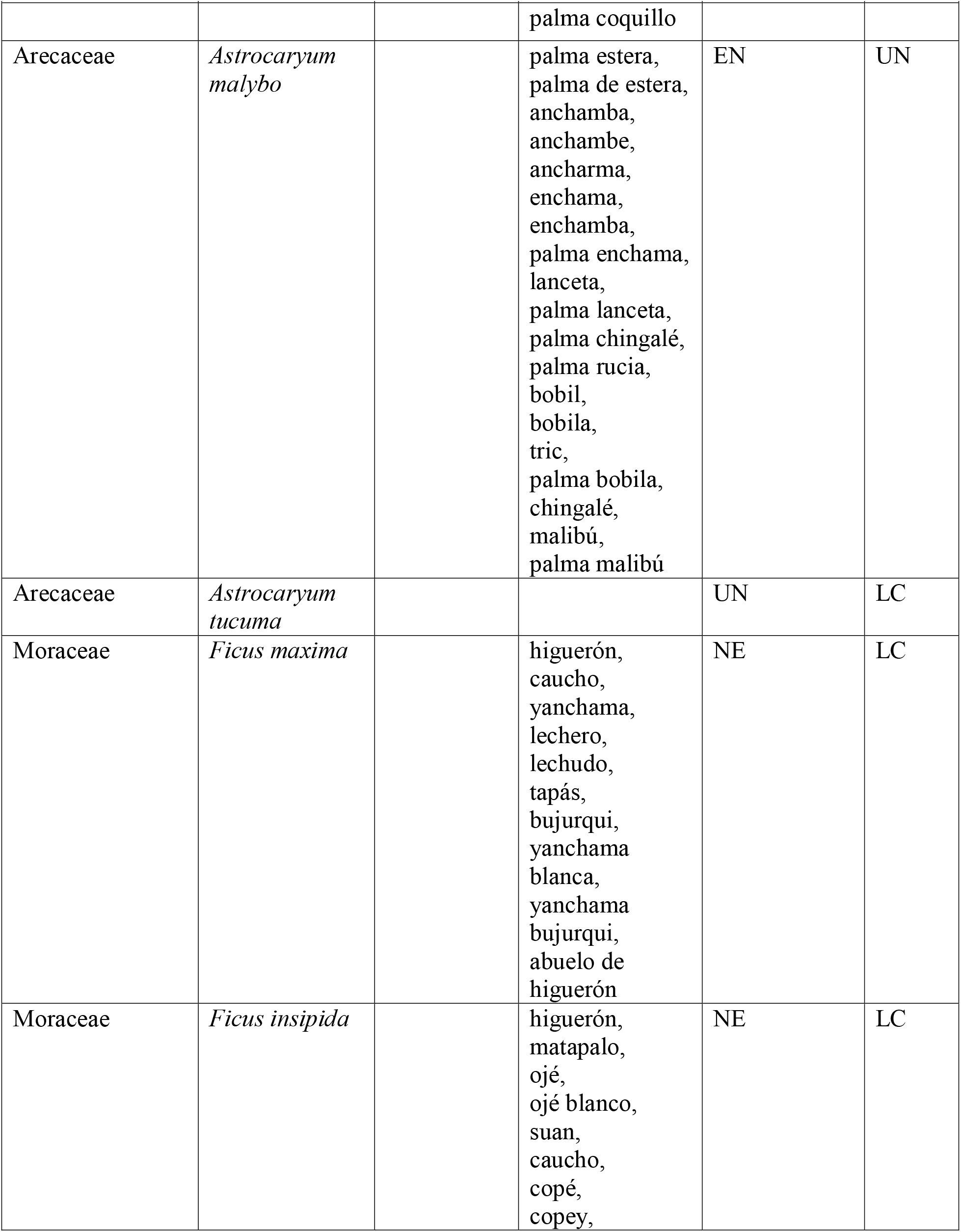
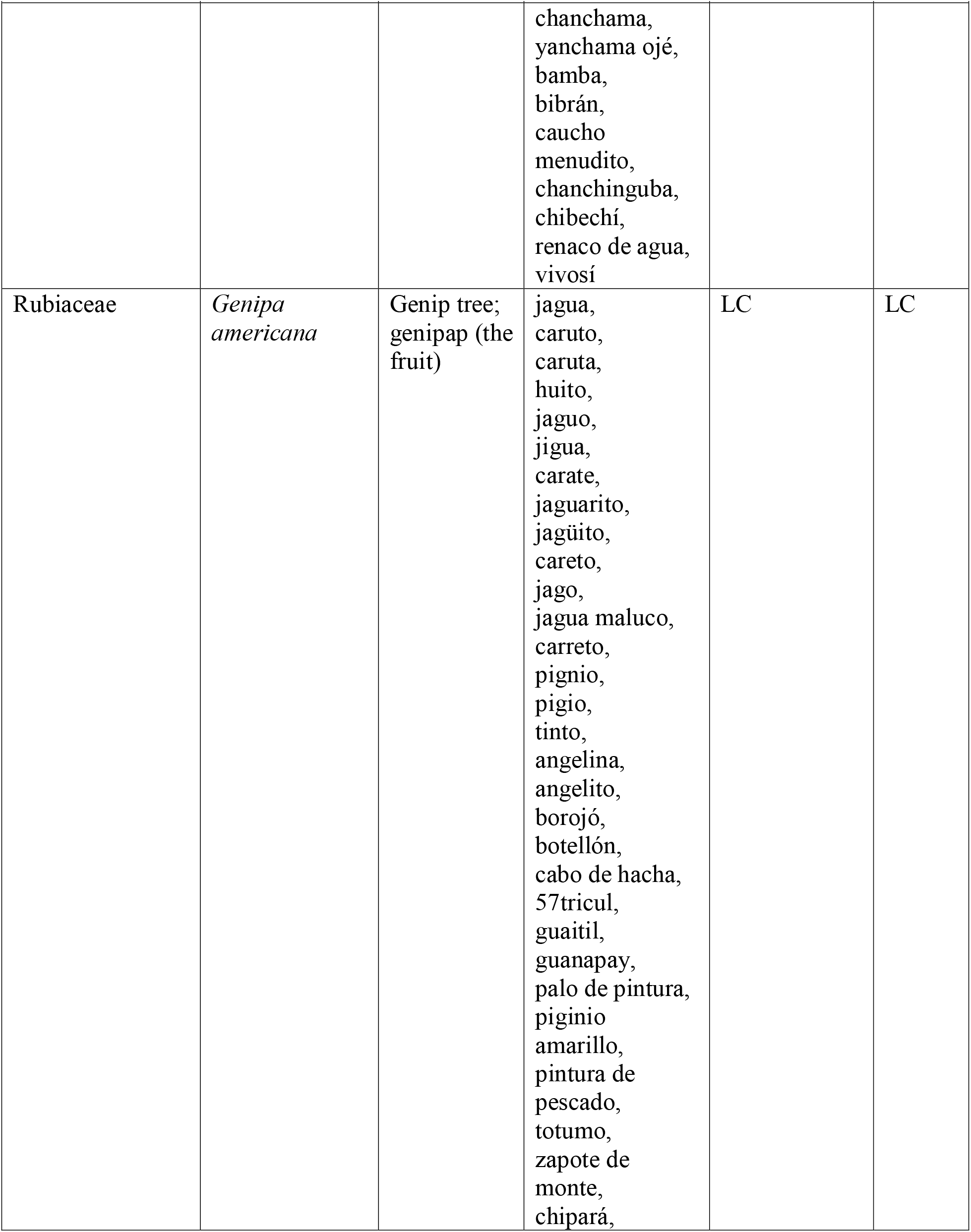
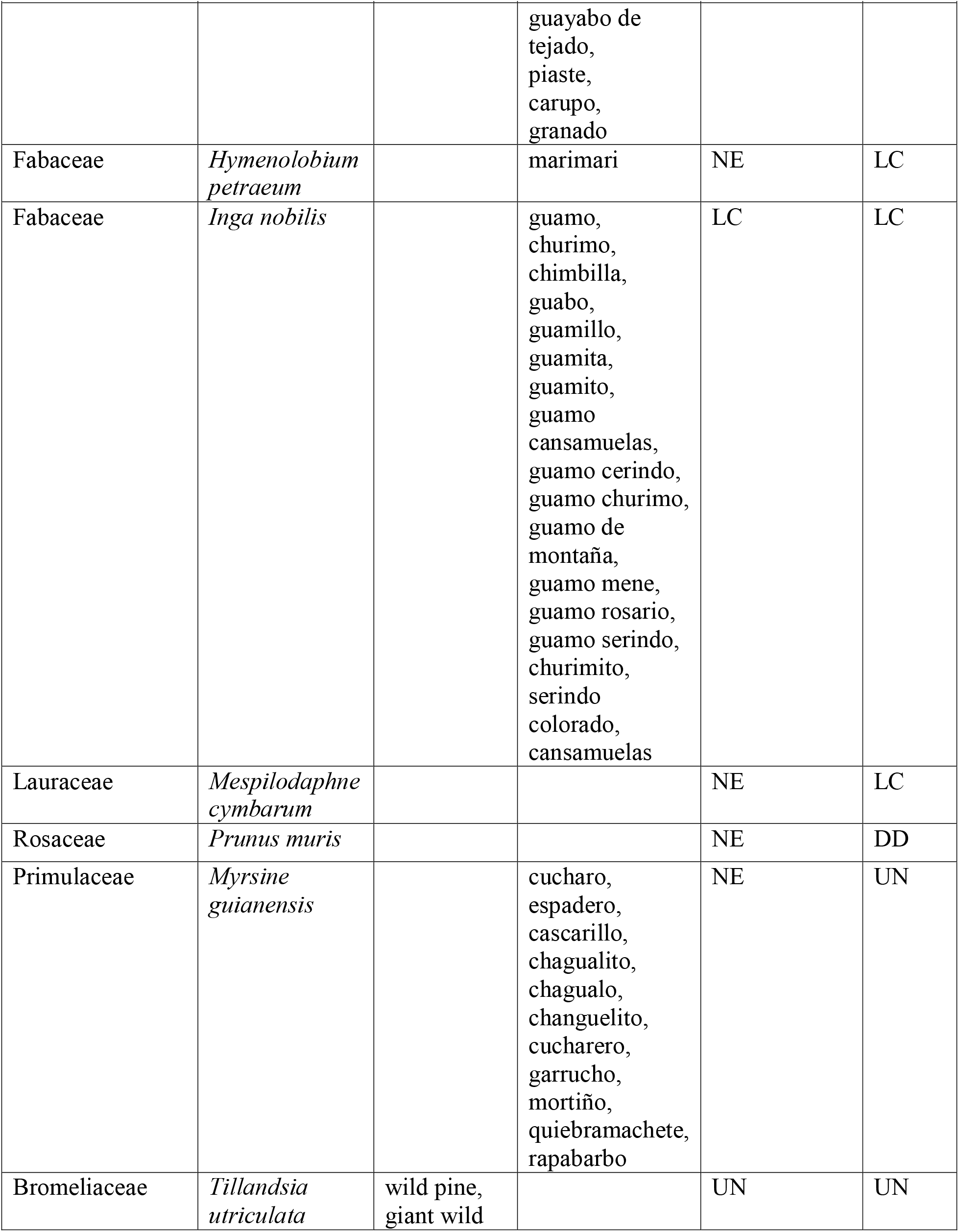
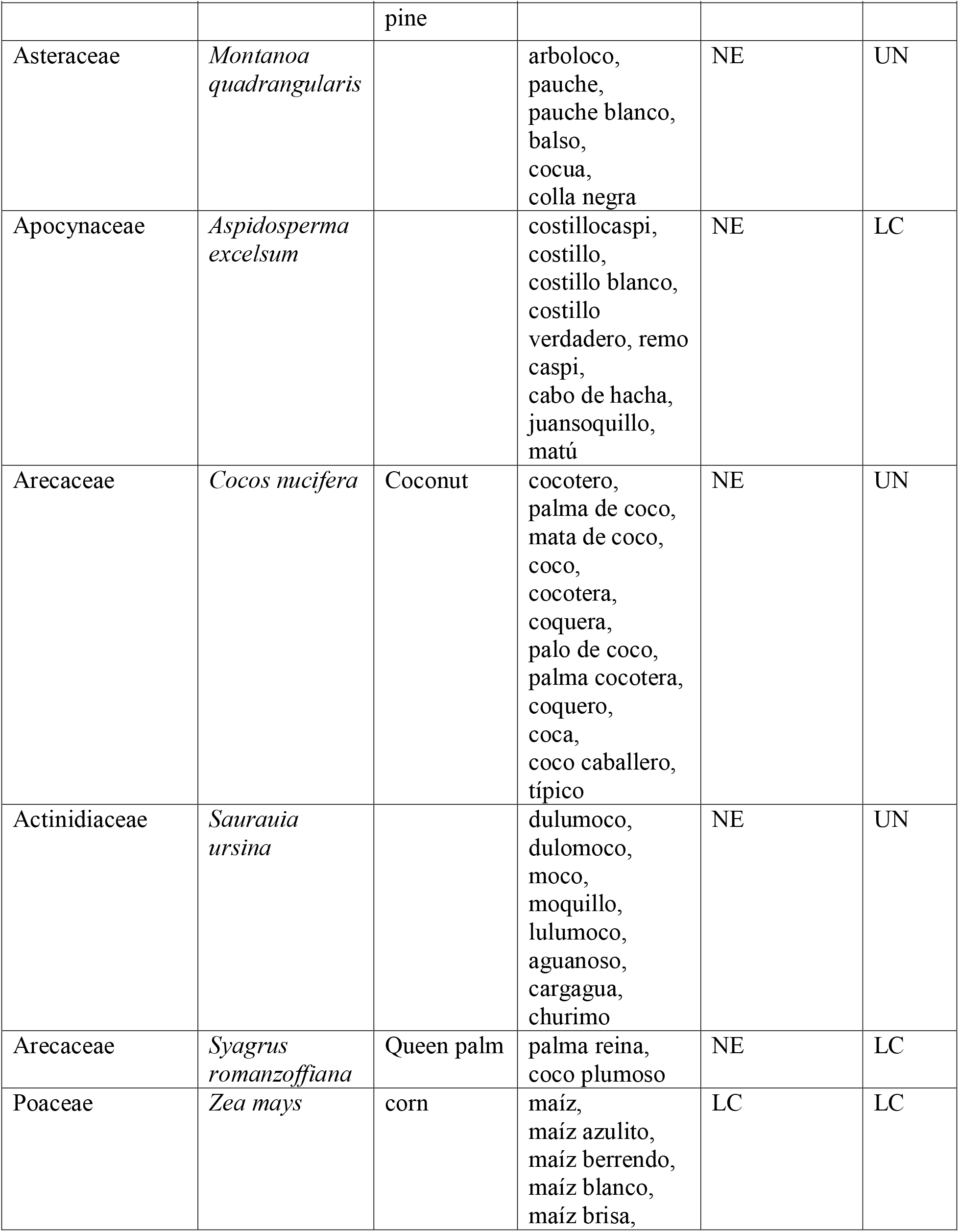
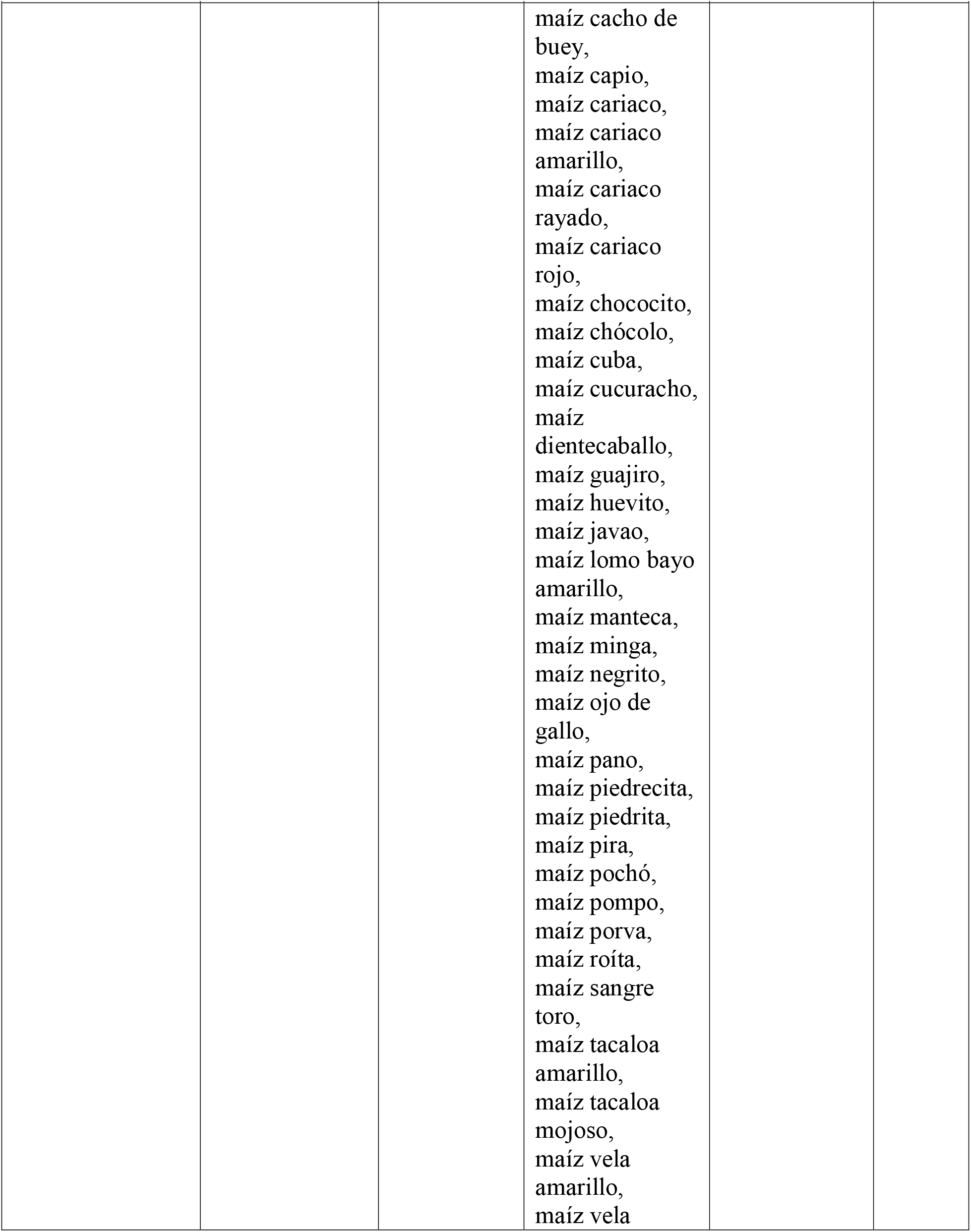
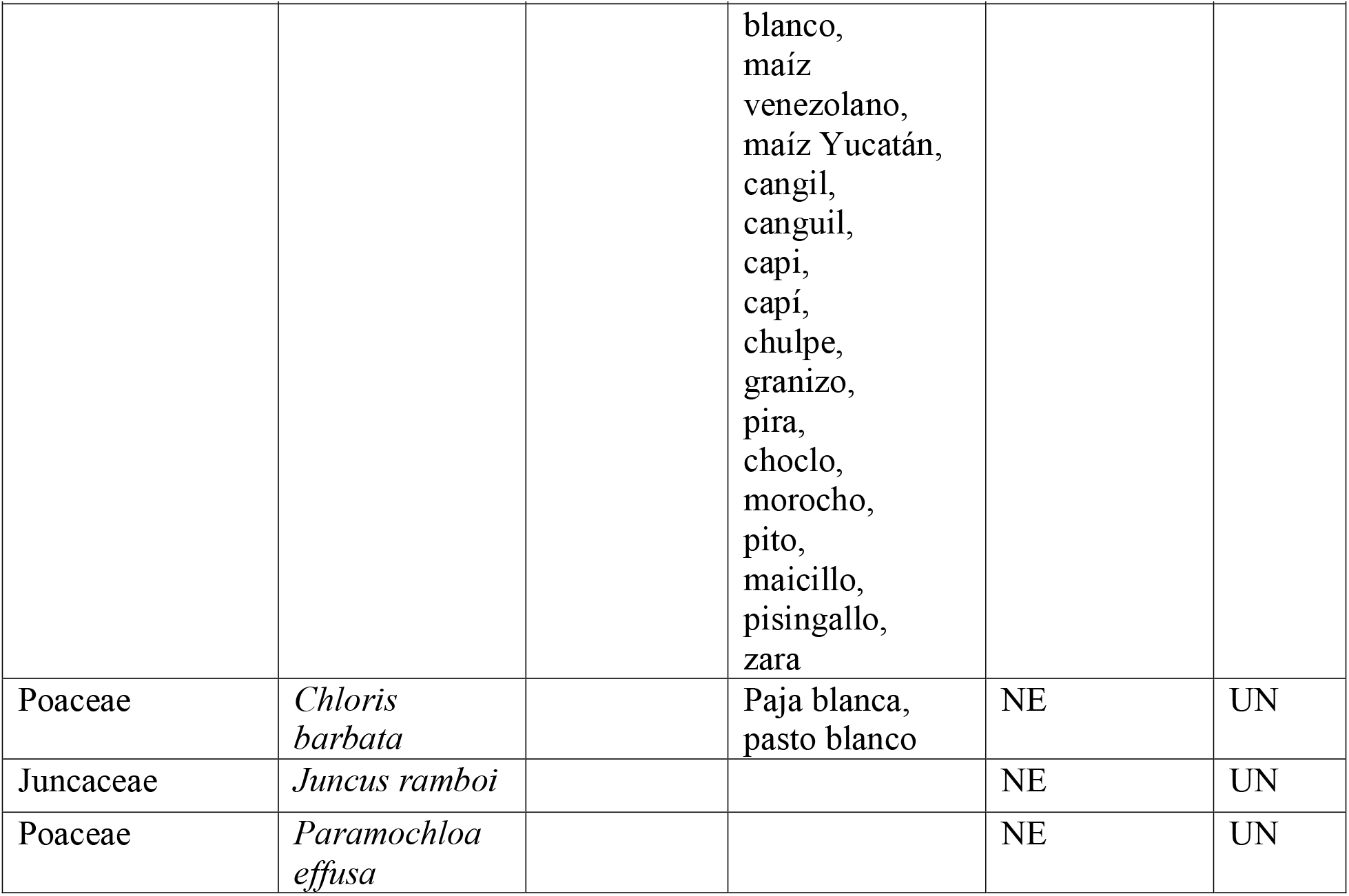
List of species used in artesanias with their scientific names, English names, and local names.

**Plot S.1.**
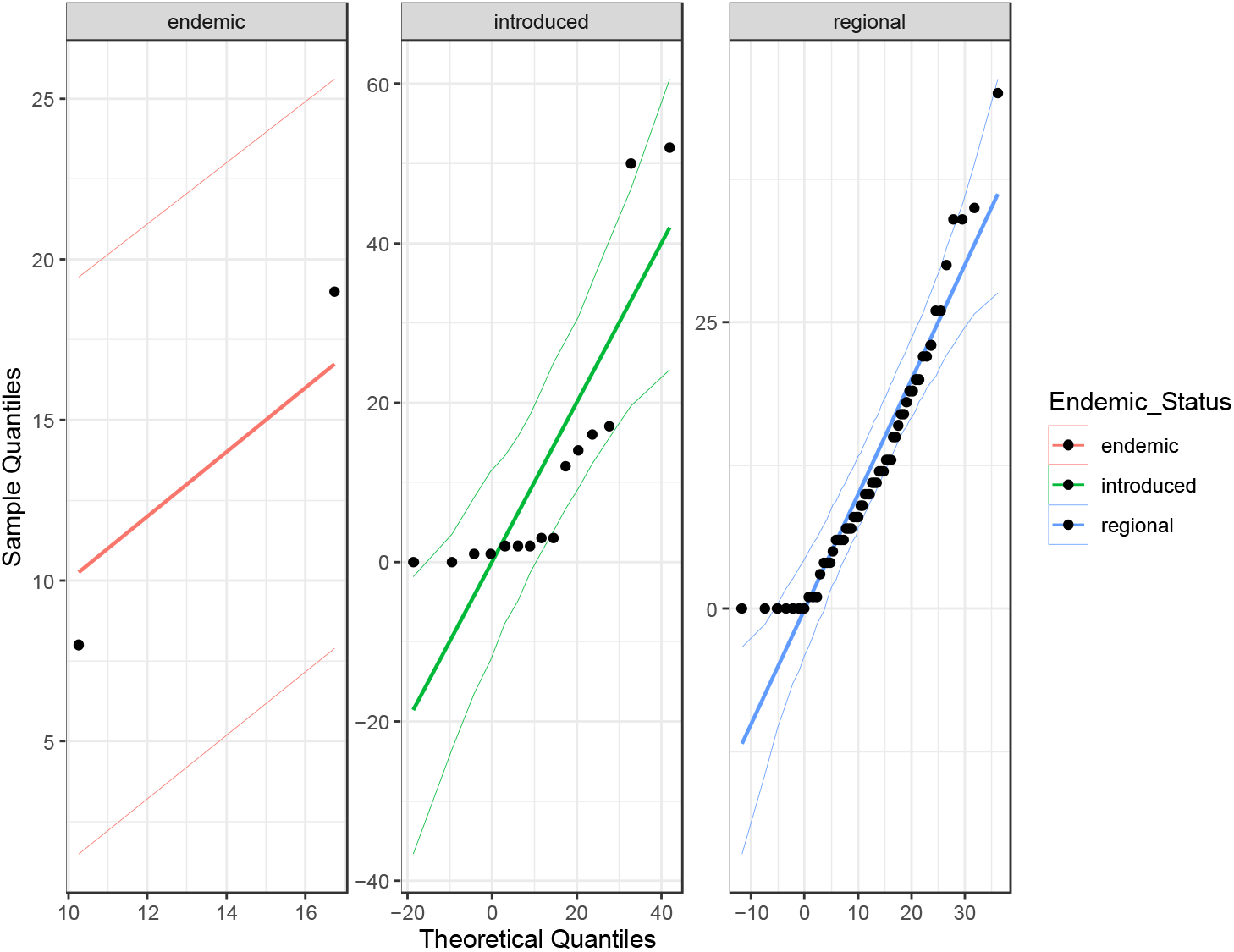
Quantile-quantile plot of number of local common names by endemic, regional, and introduced Status (Endemic_Status).

**Plot S.2.**
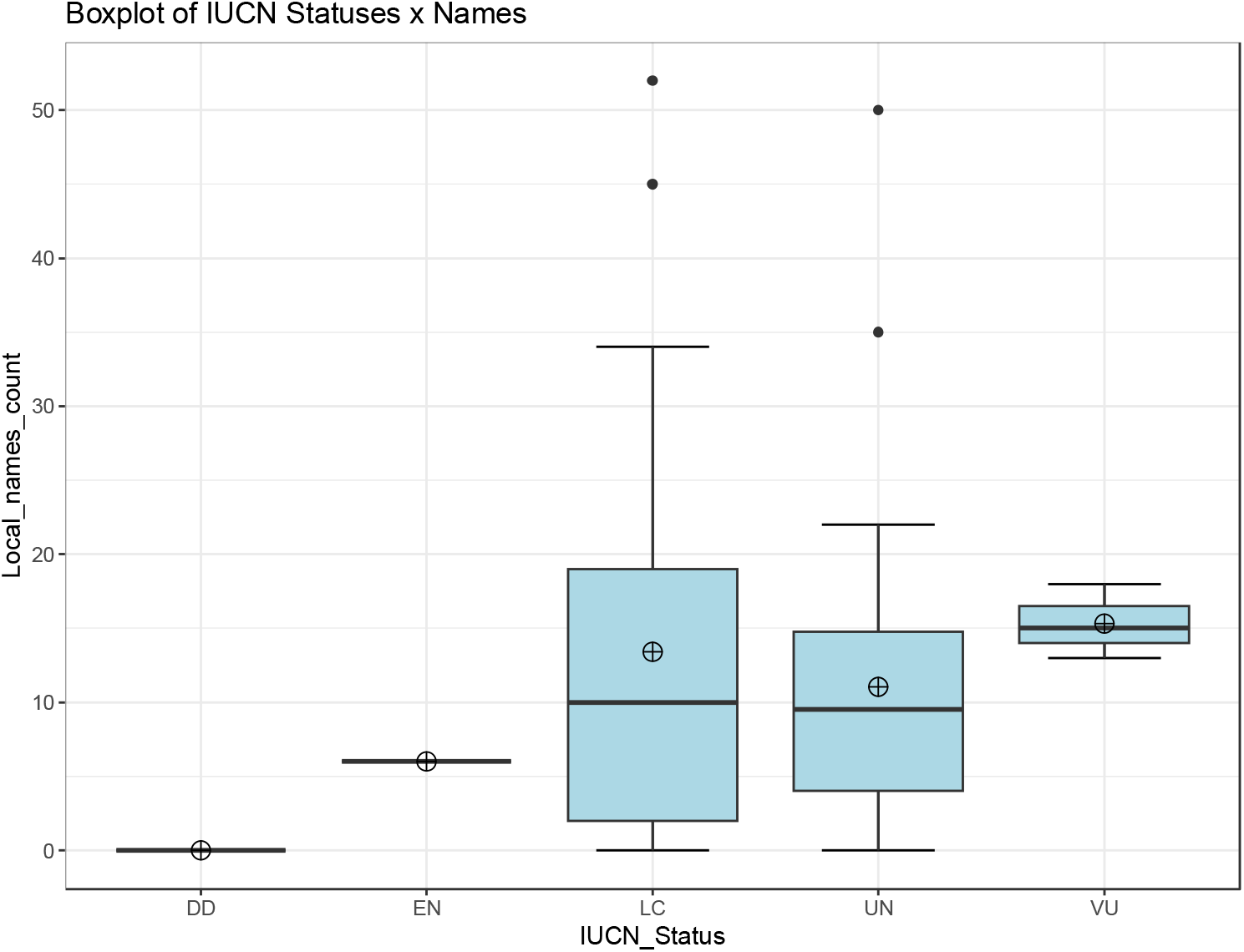
Boxplot of number of local common names (Local_names_count) by international vulnerability status (IUCN_Status).

**Plot S.3.**
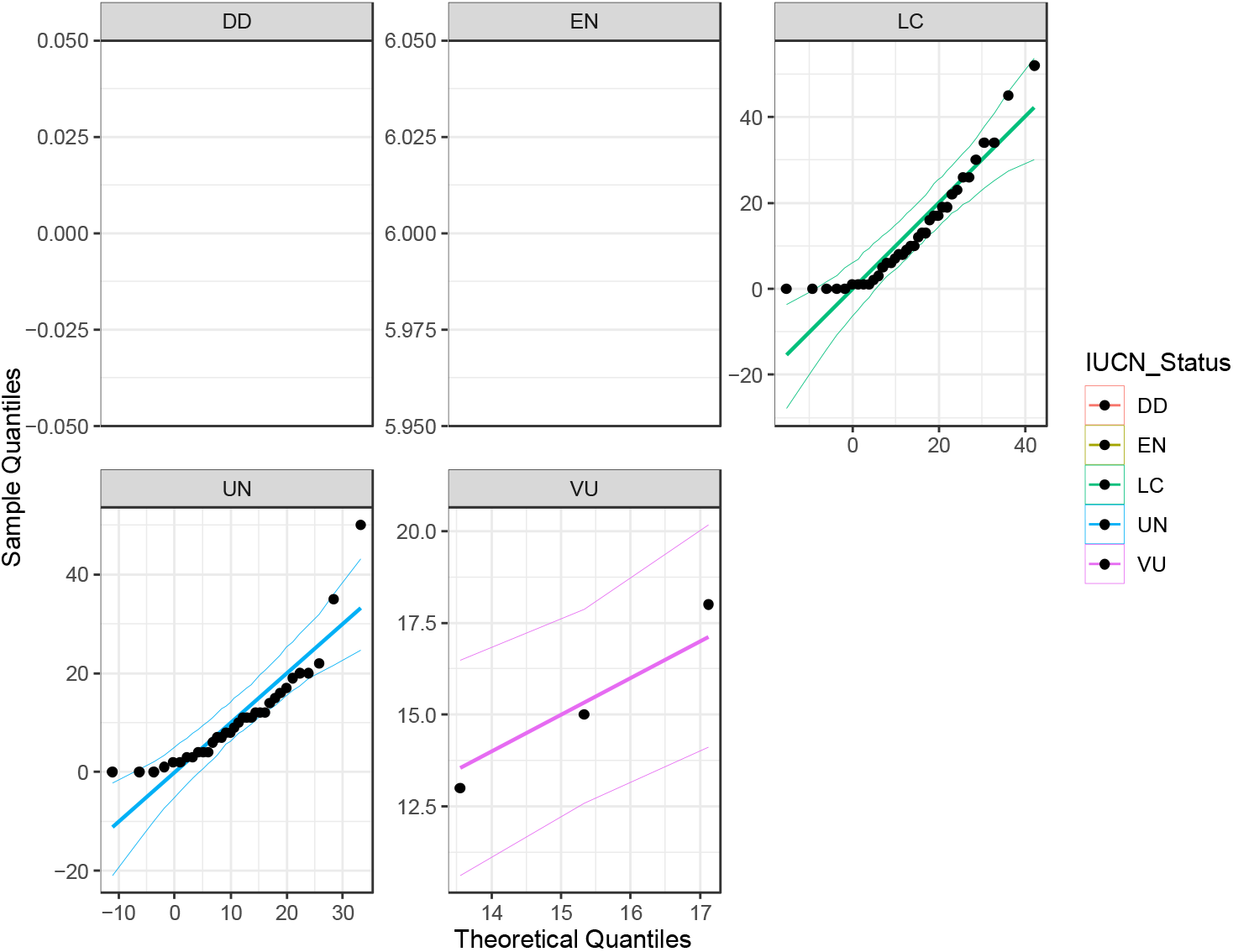
Qunatile-quantile plot of number of local common names by international vulnerability status (IUCN_Status).

**Plot S.4.**
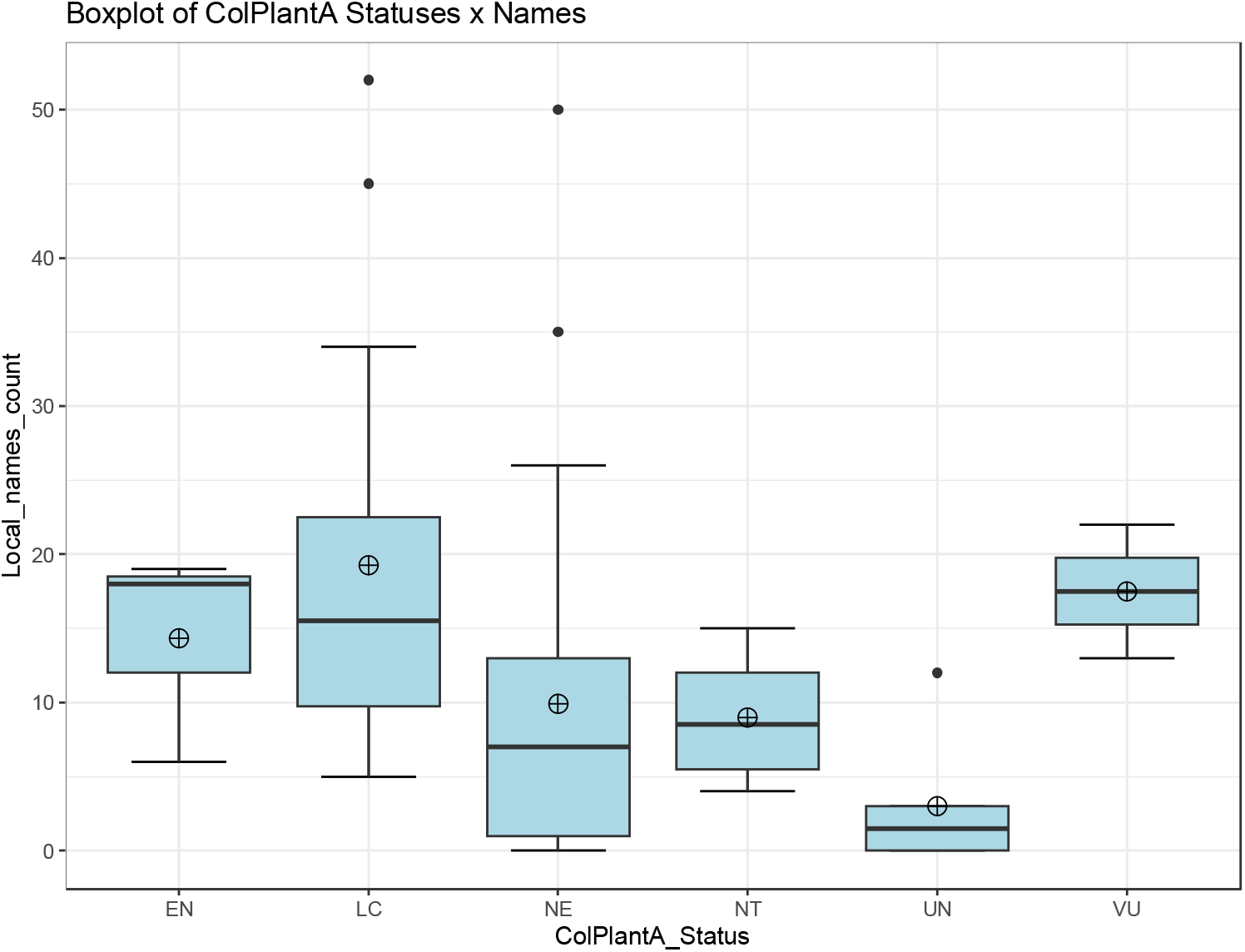
Boxplot of number of local common names (Local_names_count) by national vulnerability status (ColPlantA_Status).

**Plot S.5.**
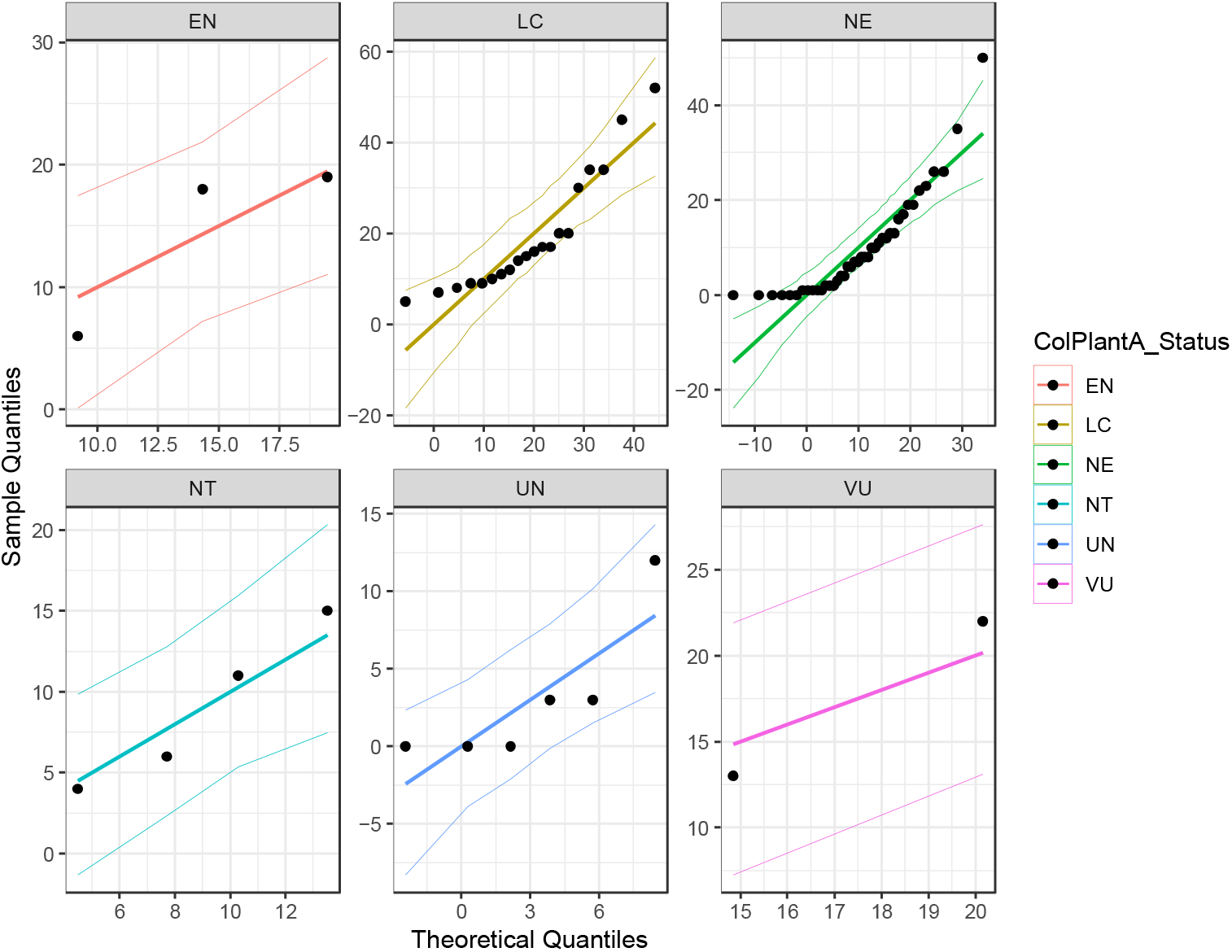
Qunatile-quantile plot of number of local common names by national vulnerability status (ColPlantA_Status).

**Plot S.6.**
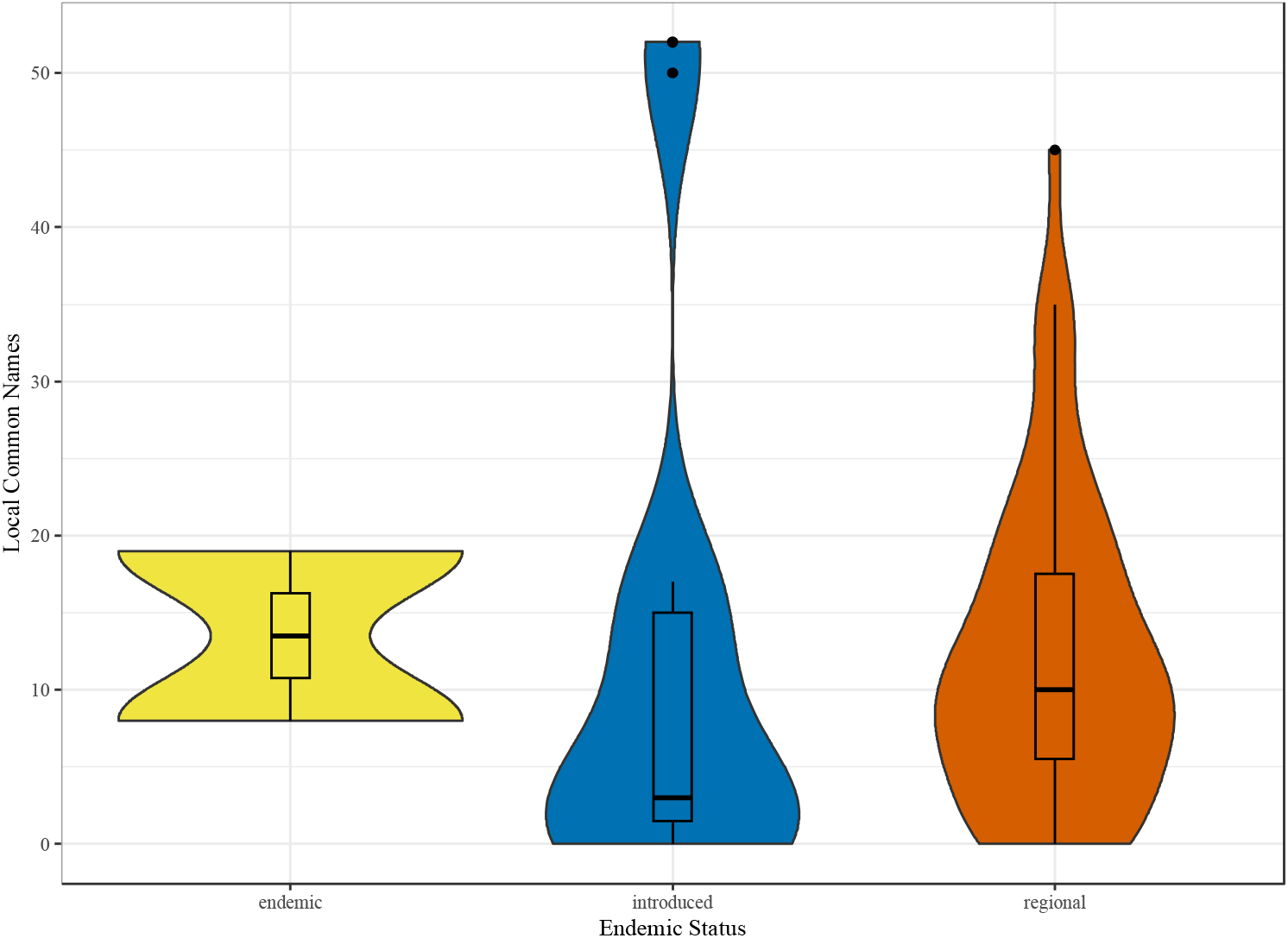
Violin plot of plant endemic status by number of local common names, with outliers included.

**Plot S.7.**
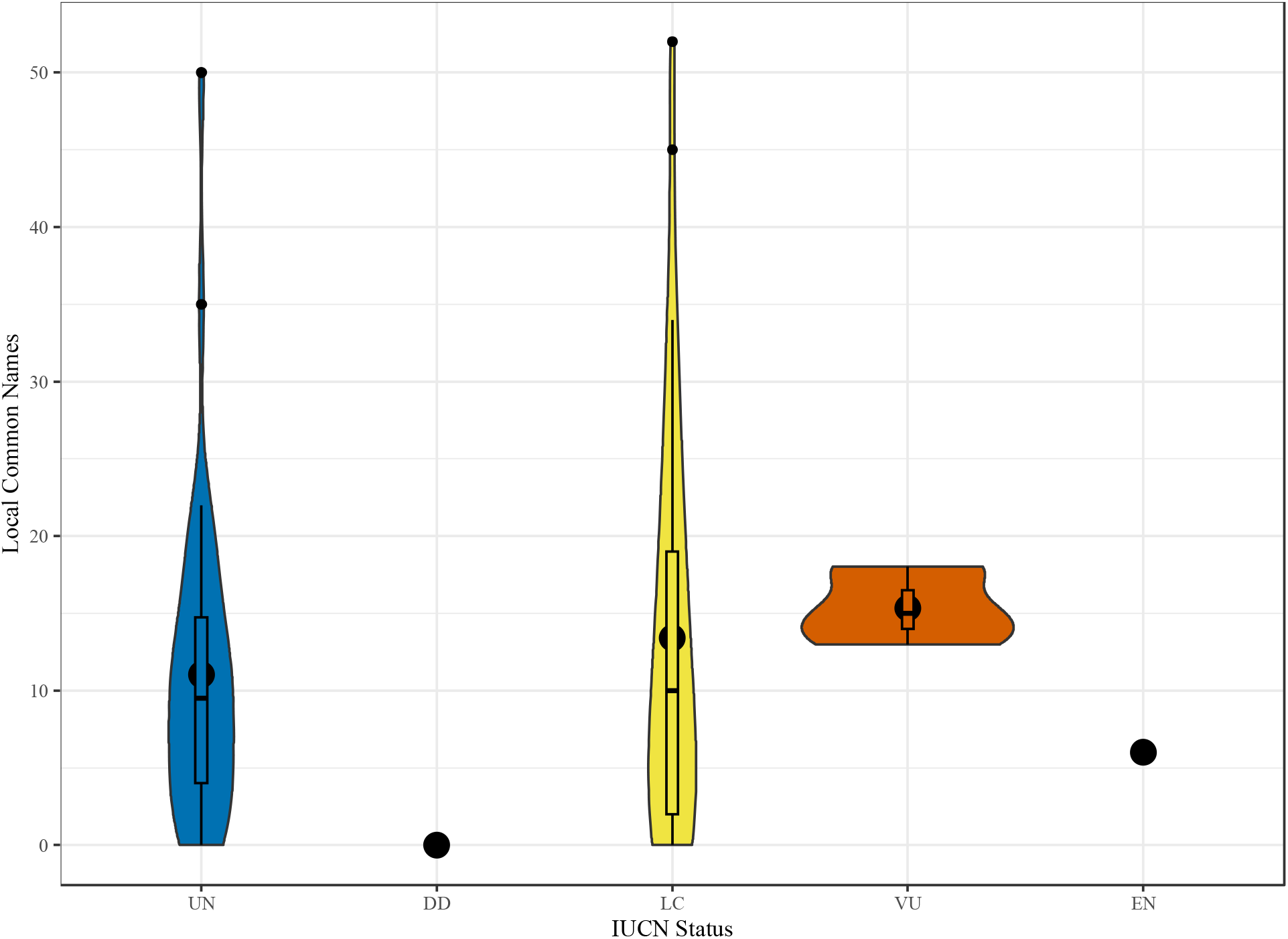
Violin plot of plant international vulnerability status (IUCN) by number of local common names, with outliers included.

**Plot S.8.**
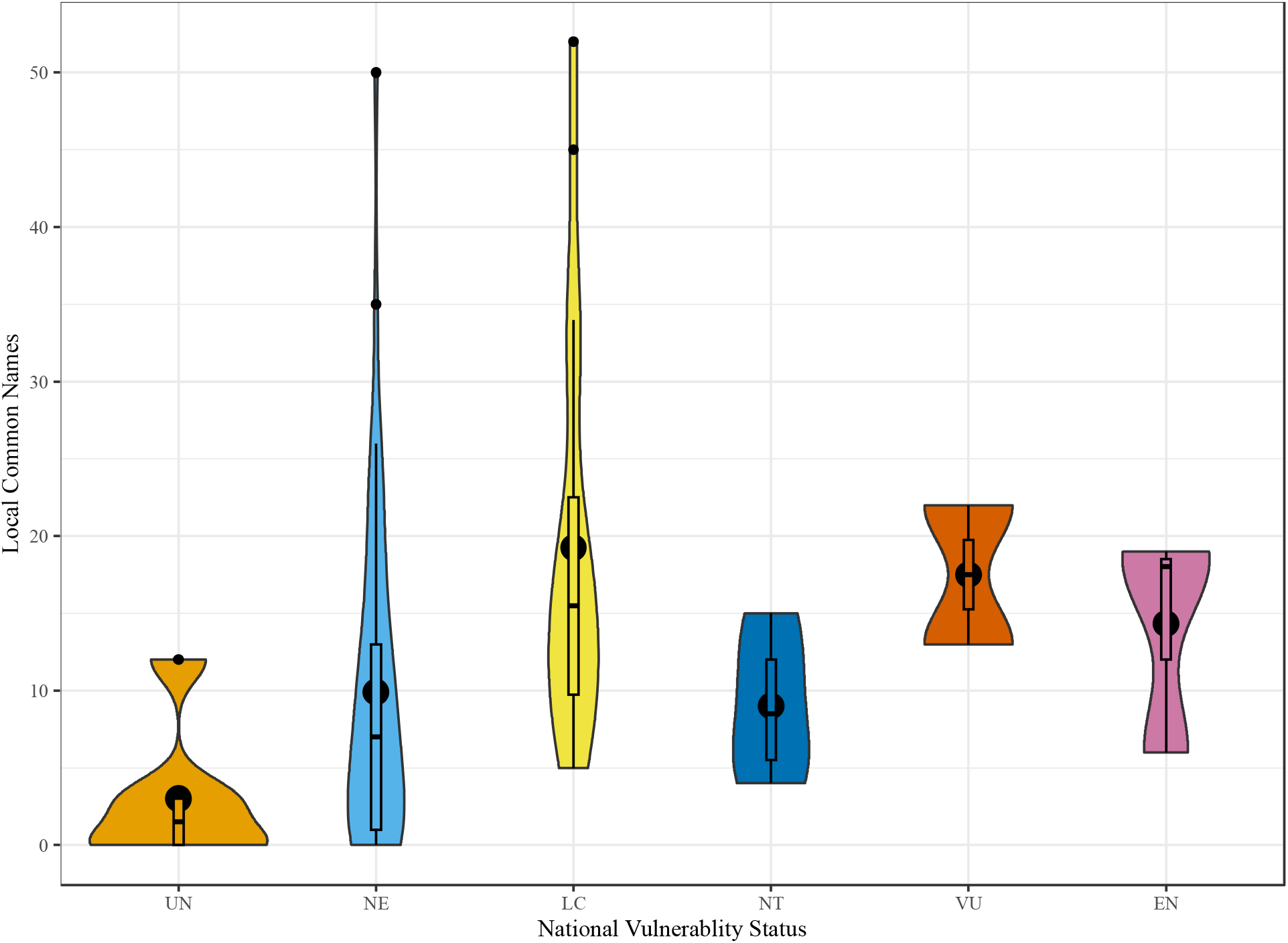
Violin plot of plant national vulnerability status (from Catálogo de Plantas y Líquenes de Colombia (Bernal et al., 2019)) by number of local common names, with outliers included.

